# Grape Expectations: Disentangling Environmental Drivers of Microbiome Establishment in Winegrowing Ecosystems

**DOI:** 10.1101/2025.05.08.652874

**Authors:** Lena Flörl, Patrik Schönenberger, Markus Rienth, Nicholas A. Bokulich

## Abstract

Microbial communities play an integral role in agricultural systems, where their composition and function are shaped by environmental factors and interactions with the plant host. In viticulture and winemaking, the resulting distinct microbial biogeographic patterns influence wine characteristics – a well-established concept termed *microbial terroir*. Yet, due to the complexity of interactions, the specific environmental drivers shaping these microbiomes remain poorly understood.

We conducted a multi-year, large-scale survey (N = 680 samples) of Swiss vineyards (N = 95 sites), following a subset (N = 12, within 2.46 km) longitudinally over 3 years. We integrate microbiome (bacterial and fungal marker-gene sequencing), untargeted metabolomics (GC-MS and LC-MS/MS), environmental monitoring, and sensory data to disentangle abiotic factors influencing community assembly and interactions in vineyards and in wine fermentations.

Results show that topography and climate collectively structure microbial communities, yet exhibit distinct influences on soil- and plant-associated microbiomes. Berry-associated fungal communities exhibit the strongest site-specific imprint, enabling machine-learning-based prediction of even subtle microclimatic differences. Climatic factors and berry chemistry exhibit an inverse relationship, selectively favoring either *Hanseniaspora* sp. or *Saccharomyces cerevisiae* – each correlating with distinct metabolite and aroma profiles. Additionally, plant stress-response metabolites were associated with shifts in microbial composition and fermentation outcomes.

By integrating multi-omics and machine learning approaches, this study enhances our understanding of microbial biogeography and its role in winegrowing. Our findings highlight the need for further research on microbial transmission, vineyard management, and implications for wine quality, offering a foundation for a more precise, science-driven approach to *terroir* expression.

**Graphical Abstract:** We sampled a total of 95 vineyards, with longitudinal sampling of 12 over three years, controlling for cultivar while capturing variations in topography, soil properties, and microclimate. Data collected in vineyards further included berry chemistry, vineyard management, and phenology. We sequenced microbial communities in both vineyard and fermentation samples, and the wines were additionally analysed with three different untargeted metabolomics approaches. Supervised and unsupervised modeling revealed spatio-temporal variability, the influence of environmental factors on microbial communities, and their impact on wine characteristics.

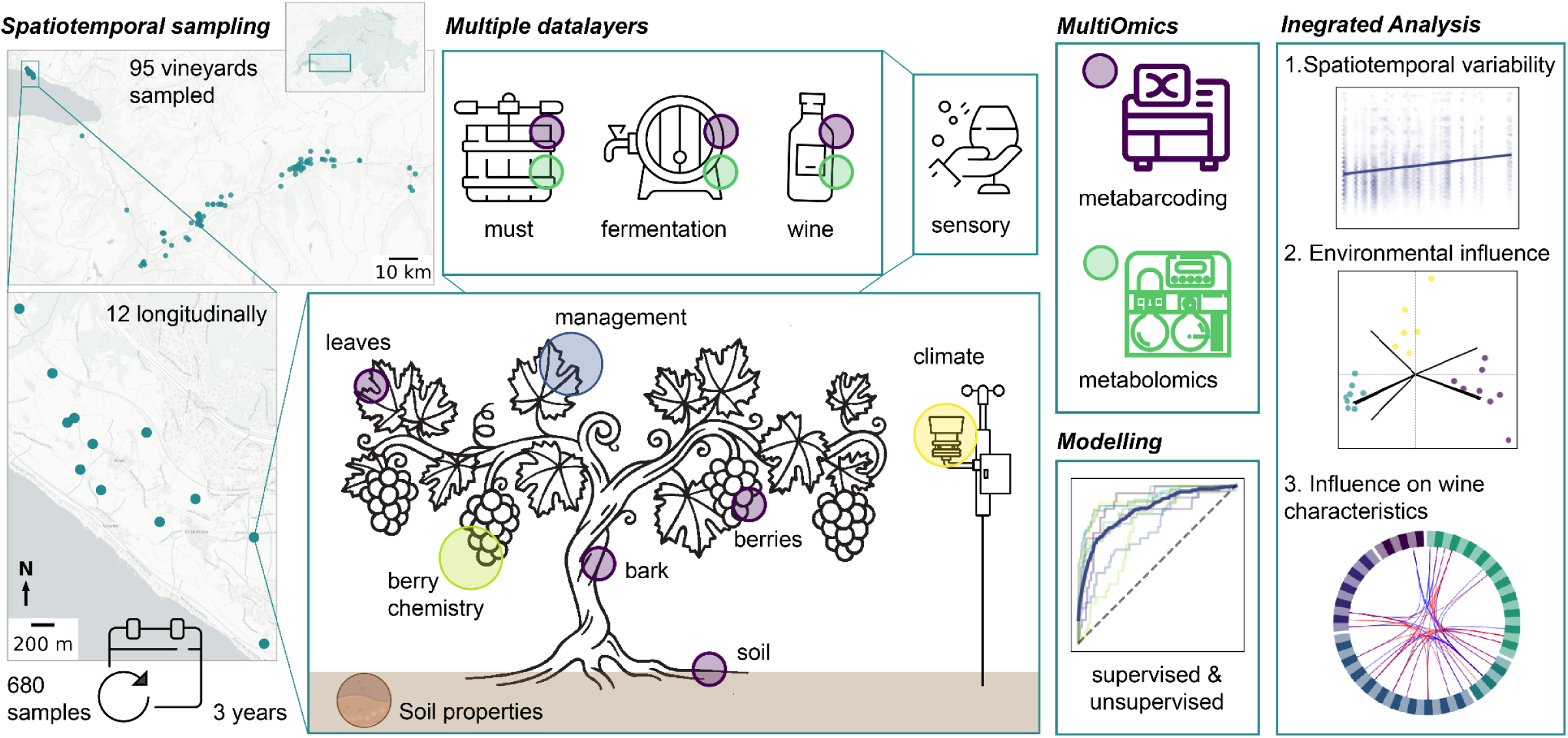

## Introduction

Microorganisms and microbial communities interact with plants as pathogens or beneficial partners, enhancing growth, nutrient uptake, stress tolerance, and disease resistance. Plant-associated microbiomes are shaped by the host genotype as well as a plethora of environmental factors, which are notoriously hard to disentangle [1]. Grapevines (*Vitis vinifera*) are a unique biogeographic model for investigating associated microbial community dynamics. As long-lived perennials cultivated across the globe in diverse environments, grapevines undergo annual recolonization of plant organs, providing an ideal system to study microbial dispersal, niche adaptation, and temporal stability. Additionally, grape berry-associated microbes can directly impact fermentation and influence resulting wine characteristics, which makes vineyard and wine microbiomes an interesting case for studying microbial dynamics and transmission in fermented foods [2]. These contributions of regionally variable microbiota to wine characteristics is termed *microbial terroir* [3], and extends our traditional understanding of *terroir* – encompassing the influence of site-specific factors including environmental variables (e.g. soil properties, climate), cultivar, rootstock, and management practices. Vineyard microbiomes are directly influenced by multiple biotic and abiotic factors, including climate and vintage effects [4], cultivar [5–7] and rootstock [8], management practices [9,10], altitude [11], soil chemistry [12], and surrounding land use [13]; and are predictive of wine quality downstream [4,14]. Yet, while numerous studies from major wine-growing regions worldwide have identified microbial patterns in the context of individual environmental influence factors and for different plant organs, a comprehensive understanding of specific contributions and underlying casualties of site specific factors on vineyard microbial communities as a whole remains elusive.

In this study, we aimed to disentangle spatial, climatic, and biotic factors influencing microbial assembly in vineyards, and their impact on resulting wine properties. We conducted the first large-scale survey of regionally varying microbiota in Swiss vineyards, sampling a total of 95 vineyards. From these, we collected longitudinal samples from 12 vineyards within a maximum distance of ∼2.5 km from each other over three consecutive years. By integrating detailed environmental data from this highly controlled setting with identical cultivars, under different microclimatic conditions and topographical features (altitude, solar exposure, etc.), while controlling for several covariates that can be challenging to control in commercial vineyards. We collected samples from the same grape variety (Chasselas) grafted on identical rootstock across distinct locations, and each year performed microvinifications. Samples were analysed with three distinct untargeted metabolomics methodologies and sensory analysis. By analyzing these diverse data layers and multi-omics datasets using machine learning and integrated analysis, we gained new insights into microbial community assembly as a function of site. This work thereby enhances our understanding of microbial contributions to *terroir* and provides fundamental insights into plant-microbiome interactions under variable environmental conditions, as well as their effects on fermentation and wine characteristics.

## Material and Methods

### Vineyard Sites: Sample and Data Collection

Samples were collected from vineyards in the Appellation d’origine contrôlée (AOC) Valais in 2023 and from 12 vineyards in Villette AOC Lavaux over 3 years (2021 to 2023). Detailed descriptions of longitudinal sampling, data collection, berry ripeness, soil chemistry and sample processing is described in the Supplementary Methods.

### Library Preparation, Sequencing and Microbiome Analysis

We used the HighALPS ultra-high throughput library preparation protocol [15] to profile bacterial (V4 region of the 16S rRNA gene using 515F and 806R primers [16,17]) and fungal (BITS and B58S3 primers [18]) communities. Subsequent microbiome analysis was conducted using QIIME 2 (version 2024.10) [19]. Briefly, amplicon sequence variants (ASVs) were generated by denoising with q2-dada2 [20], taxonomically classified with naive Bayes using the q2-feature-classifier plugin [21,22] trained against the SILVA or UNITE databases [23–25]. Diversity analysis was performed using the vegan package (also via q2-diversity) and q2-kmerizer [26,27] plugins for QIIME 2. Details of library preparation, bioinformatics, and statistical analysis are described in the Supplementary Methods.

### Microvinifications and Analysis of Resulting Wines

Microvinifications were performed every year using berries from each individual plot. The resulting wines were assessed by a trained sensory panel and analysed with untargeted metabolomics, namely headspace solid-phase microextraction gas chromatography–mass spectrometry (HS-SPME-GC-MS) and liquid chromatography-mass spectrometry (LC-MS). For details, see the Supplementary Methods.

## Results

We aimed to identify the environmental factors shaping microbial community assembly in vineyards, while controlling for biological variables that often confound microbial biogeography research in commercial agriculture. Here, we focus on an existing vineyard network in the Appellation d’origine contrôlée (AOC) Lavaux that has been deeply characterized for phenology, climate, pedology, and *terroir* expression [28], planted exclusively to Chasselas cultivar and 3309C rootstock to control for biotic factors influencing microbial community assembly in grapevines [5,7]. Integrating microbiome analysis, metabolome analysis, and continuous environmental monitoring in this vineyard network thus enables an exquisite opportunity to study how spatial, environmental, climatic, and microbial factors interact with each other and with wine quality parameters downstream.

### Vineyards Within AOC Lavaux Exhibit a Gradient of Pedoclimatic Characteristics

First, we collected samples and data over three years from 12 distinct vineyards planted with Chasselas in the AOC Lavaux subregion “Villette”, situated within a radius of 2.46 km. The plots were chosen to represent distinct pedoclimatic characteristics present within the AOC, which are known to contribute to the specific environmental context of a vineyard such as inclination, altitude, and solar radiation. Additionally, we continuously measured temperature and relative humidity at each site and assessed soil characteristics, phenology, management practices and other agronomic information (e.g., spread of mildew, or hail damage). From the climate measurements we calculated the daily median values and coefficient of variation to assess fluctuations per vineyard over the growing season. This revealed that 2021 was substantially more humid and cool (median relative humidity (RH): 71.98 %, median temperature: 18.51°C) than 2022 and 2023 (median RH: 63.39% and 65.71%, temperature: 20.81°C and 20.58°C, respectively). This variability is further reflected in the berry chemistry composition, e.g., substantially higher malate concentrations in 2021 due to cooler temperatures and consequently reduced malate respiration during ripening. In contrast, the average berry composition in 2022 and 2023 remained relatively consistent. This reflects the large impact of annual variations on berry development and chemical composition, which depend on the respective site-specific and climatic conditions (see Fig. 1).

**Figure 1.**
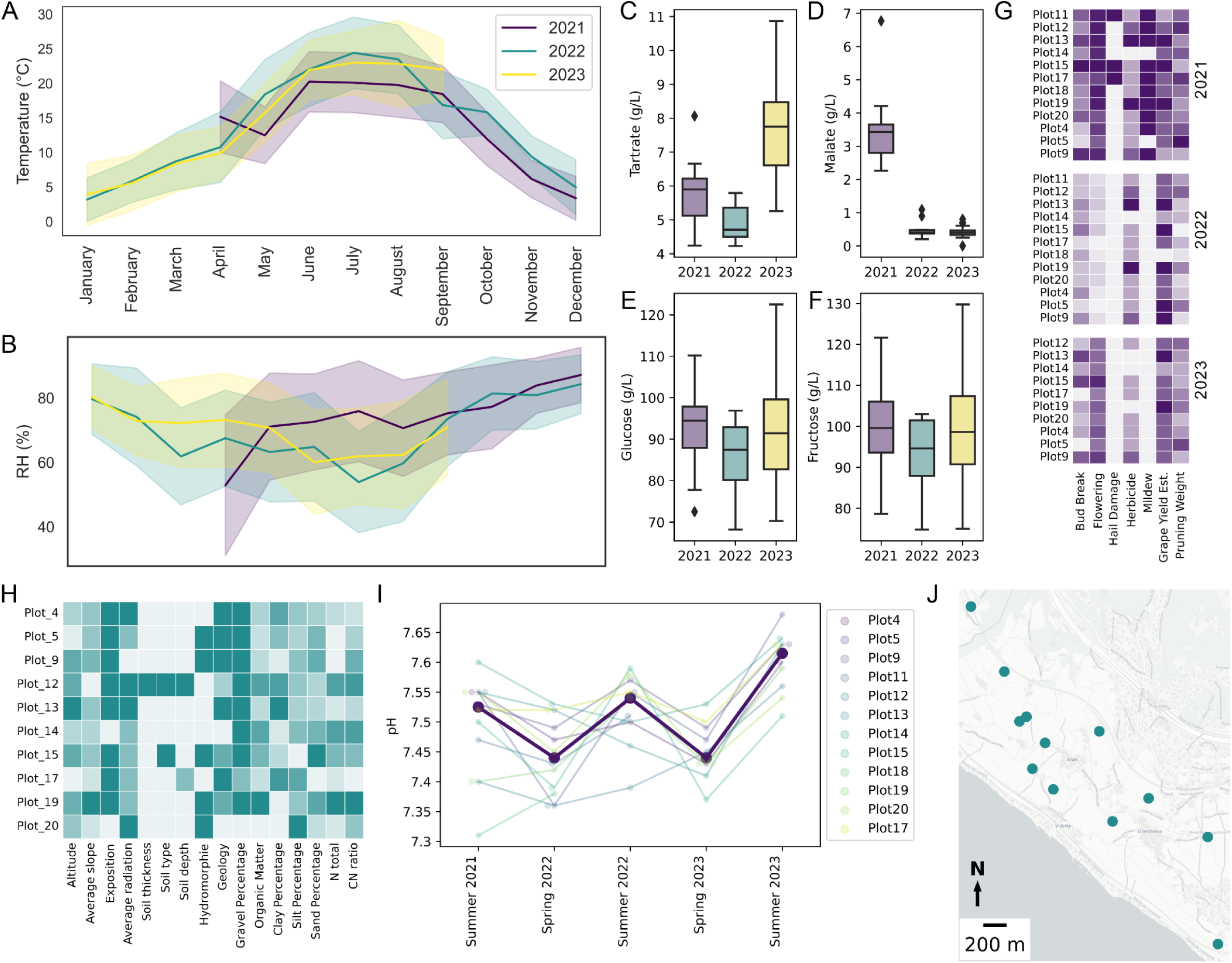
Characterization of the 12 vineyards sampled in AOC Lavaux over three years, illustrating climatic variation (A: temperature, B: relative humidity (RH)) and berry chemistry (C: tartrate, D: malate, E: glucose, F: fructose). Additionally, key agronomic and phenological metrics were assessed annually (G), including bud break and flowering time, hail damage, mildew occurrence, grape yield estimates, and pruning weight as a proxy for plant vigor. (H) All vineyards exhibited distinct pedoclimatic properties, including altitude, slope, solar exposure, radiation, soil type and depth, hydromorphy, geology, gravel percentage, organic matter and nitrogen content, and soil texture. (I) Soil pH was also measured throughout the years, revealing seasonal fluctuations. (J) Shows the locations of the sampled vineyards within the AOC.

### Vineyard Microbiomes Vary in Their Microbial Richness and Composition, and Exhibit Distinct Spatio-temporal Patterns

In total we collected 680 samples, consisting of 441 berry samples, 98 samples from subsequent microvinifications (laboratory-scale fermentations conducted from these berries, see Methods), 55 soil samples, 53 leaf samples and 33 bark samples. Samples were analyzed using marker-gene amplicon sequencing for bacterial (16S rRNA gene) and fungal (ITS) communities, yielding 18’795’287 reads for fungi and 3’306’142 reads for bacteria. Due to low sequencing depth for bacterial communities in most grape and leaf samples (mean read count <700), these samples were excluded from the beta diversity analysis. The different vineyard sample types exhibit distinct differences in alpha diversity (Kruskal Wallis for Fungi: p_Richness_ = 2.30e-39, p_Eveness_ = 1.56e-46, p_Shannon Entropy_ = 1.02e-42; Bacteria: p_Richness_ = 8.48e-19, p_Eveness_ = 2.15e-10, p_Shannon Entropy_ = 2.73e-17) with a gradient in richness from soil to bark to leaves to berries; and beta diversity, with soil being most distinct from annually newly formed plant organs of leaves and berries. The latter also showed substantial overlap, particularly for the presence of fungi (see Supplementary Figure 1).

Next, we analyzed spatio-temporal variation in fungal and bacterial alpha and beta diversity of individual sample types within and between vineyard sites. In soils, significant spatiotemporal variation was observed in bacterial community evenness and abundance (Supplementary Table 1), but not for fungi (Supplementary Table 2). Yet the composition of both fungal and bacterial communities exhibited significant variation across different vineyards and sampling time points (spring, summer) (Table 1, Table 2). Notably, the phylogenetic structure of fungal communities significantly varied between timepoints and years, even when comparing samples from the same vineyards. The vineyard had the overall strongest influence on community composition, accounting for 27.3% to 38.7% of the variation in bacterial communities and 30.5% to 39.8% in fungal communities, with considering abundance overall increasing the explained variation. However, no significant distance-decay relationship was observed (Supplementary Table 3), indicating that spatial proximity was not associated with microbial community similarity in these soils. Similarly to soil, fungal communities of bark significantly varied between vineyards, with up to 45% of variance explained by site, but without physical proximity of vineyards significantly contributing to this variation (Supplementary Table 4). These communities appear to be temporally stable as no significant differences were detected between years (Table 1). In contrast, bacterial communities did not significantly vary by location but their phylogenetic composition differed substantially between years.

**Table 1.** Permutational analysis of variance (PERMANOVA) of the various beta diversity metrics (Bray Curtis, Jaccard, Bray Curtis of kmers, Jaccard of kmers) of fungal communities for the different vineyard sample types (berries, soil, bark and leaves).

|  | Bray Curtis |  | Jaccard |  | Kmer: Bray Curtis |  | Kmer: Jaccard |  |
| --- | --- | --- | --- | --- | --- | --- | --- | --- |
|  | R2 | p-value* | R2 | p-value* | R2 | p-value* | R2 | p-value* |
| **Berries** |  |  |  |  |  |  |  |  |
| Plot ID | 0.106 | **0.001** | 0.051 | **0.001** | 0.161 | **0.001** | 0.078 | **0.001** |
| Year | 0.015 | **0.001** | 0.012 | **0.001** | 0.027 | **0.001** | 0.025 | **0.001** |
| Time point | 0.028 | **0.001** | 0.006 | **0.001** | 0.034 | **0.001** | 0.013 | **0.001** |
| Plot ID:Year | 0.043 | **0.003** | 0.033 | **0.001** | 0.057 | **0.001** | 0.038 | **0.001** |
| Plot ID:Time point | 0.044 | **0.006** | 0.026 | 0.993 | 0.051 | **0.001** | 0.032 | **0.004** |
| Year:Time point | 0.018 | **0.001** | 0.003 | **0.044** | 0.022 | **0.001** | 0.004 | **0.022** |
| Residuals | 0.746 | NaN | 0.869 | NaN | 0.647 | NaN | 0.811 | NaN |
| **Soil** |  |  |  |  |  |  |  |  |
| Plot ID | 0.340 | **0.001** | 0.305 | **0.001** | 0.398 | **0.001** | 0.375 | **0.001** |
| Year | 0.028 | **0.002** | 0.027 | **0.001** | 0.031 | **0.001** | 0.031 | **0.001** |
| Time point | 0.047 | **0.001** | 0.028 | **0.001** | 0.064 | **0.001** | 0.035 | **0.001** |
| Plot ID:Year | 0.214 | 0.245 | 0.239 | 0.105 | 0.195 | 0.043 | 0.216 | **0.047** |
| Plot ID:Time point | 0.205 | 0.064 | 0.213 | 0.286 | 0.180 | **0.033** | 0.182 | 0.368 |
| Year:Time point | 0.017 | 0.702 | 0.021 | 0.437 | 0.016 | 0.262 | 0.018 | 0.421 |
| Residuals | 0.148 | NaN | 0.167 | NaN | 0.116 | NaN | 0.142 | NaN |
| **Bark\*\*** |  |  |  |  |  |  |  |  |
| Plot ID | 0.433 | **0.049** | 0.400 | **0.015** | 0.450 | **0.021** | 0.424 | **0.004** |
| Year | 0.026 | 0.855 | 0.034 | 0.334 | 0.026 | 0.680 | 0.033 | 0.344 |
| Plot ID:Year | 0.311 | 0.727 | 0.334 | 0.450 | 0.305 | 0.590 | 0.320 | 0.462 |
| Residuals | 0.230 | NaN | 0.232 | NaN | 0.219 | NaN | 0.222 | NaN |
| **Leaves\*\*** |  |  |  |  |  |  |  |  |
| Plot ID | 0.348 | 0.318 | 0.354 | 0.887 | 0.346 | 0.201 | 0.339 | 0.996 |
| Year | 0.082 | **0.016** | 0.043 | 0.179 | 0.121 | **0.002** | 0.051 | 0.087 |
| Plot ID:Year | 0.384 | 0.204 | 0.375 | 0.626 | 0.372 | 0.145 | 0.362 | 0.957 |
| Residuals | 0.186 | NaN | 0.228 | NaN | 0.161 | NaN | 0.248 | NaN |

\* p-values < 0.05 are considered significant and highlighted in bold
\*\* Note: bark and leaves were only sampled at one timepoint, so this factor is excluded in these tests.

**Table 2.** Permutational analysis of variance (PERMANOVA) of the various beta diversity metrics (Bray Curtis, Jaccard, unweighted UniFrac, weighted UniFrac, Bray Curtis of kmers, Jaccard of kmers) of bacterial communities for the different vineyard sample types (berries, soil, bark and leaves).

|  | Bray Curtis |  | Jaccard |  | unweighted UniFrac |  | weighted UniFrac |  | Kmer: Bray Curtis |  | Kmer: Jaccard |  |
| --- | --- | --- | --- | --- | --- | --- | --- | --- | --- | --- | --- | --- |
|  | R2 | p-value* | R2 | p-value* | R2 | p-value* | R2 | p-value* | R2 | p-value* | R2 | p-value* |
| **Soil** |  |  |  |  |  |  |  |  |  |  |  |  |
| Plot ID | 0.387 | **0.001** | 0.282 | **0.001** | 0.273 | **0.003** | 0.366 | **0.013** | 0.379 | **0.012** | 0.277 | **0.001** |
| Time point | 0.055 | **0.001** | 0.033 | **0.002** | 0.043 | **0.001** | 0.069 | **0.005** | 0.060 | **0.007** | 0.038 | **0.001** |
| Year | 0.022 | 0.203 | 0.025 | 0.019 | 0.031 | **0.020** | 0.036 | 0.082 | 0.027 | 0.188 | 0.030 | **0.013** |
| Plot ID : Time point | 0.186 | 0.780 | 0.216 | 0.939 | 0.207 | 0.961 | 0.175 | 0.845 | 0.171 | 0.944 | 0.211 | 0.976 |
| Plot ID : Year | 0.131 | 0.998 | 0.195 | 0.949 | 0.195 | 0.868 | 0.122 | 0.987 | 0.120 | 0.998 | 0.191 | 0.978 |
| Time point : Year | 0.015 | 0.763 | 0.020 | 0.651 | 0.021 | 0.457 | 0.014 | 0.683 | 0.014 | 0.803 | 0.019 | 0.802 |
| Residuals | 0.204 | NaN | 0.229 | NaN | 0.231 | NaN | 0.219 | NaN | 0.229 | NaN | 0.234 | NaN |
| **Bark** |  |  |  |  |  |  |  |  |  |  |  |  |
| Plot ID | 0.420 | 0.390 | 0.437 | 0.157 | 0.434 | 0.212 | 0.387 | 0.220 | 0.415 | 0.387 | 0.435 | 0.304 |
| Year | 0.057 | 0.100 | 0.045 | 0.186 | 0.050 | 0.127 | 0.101 | **0.008** | 0.061 | 0.124 | 0.054 | **0.043** |
| Plot ID : Year | 0.364 | 0.238 | 0.353 | 0.138 | 0.362 | 0.165 | 0.407 | 0.086 | 0.373 | 0.262 | 0.345 | 0.367 |
| Residuals | 0.158 | NaN | 0.165 | NaN | 0.154 | NaN | 0.104 | NaN | 0.152 | NaN | 0.166 | NaN |

\* p-values < 0.05 are considered significant and highlighted in bold

Contrary to bark, the fungal communities on berries and leaves were less temporally stable and varied significantly between years (Table 1). Fungal communities of berries showed to be most strongly affected by spatio-temporal factors. While the evenness is significantly different between the vineyards, the number of observed taxa also varied by time point and year (Supplementary Table 1). Fungal community composition varied substantially between vineyard, ripeness stages, and years (Table 1). When accounting for abundance, the explained variance nearly doubled. Interestingly, while these effects are robust and also significant when considered in interaction with each other (e.g. for plot and year, or year and time point), the overall explained variance is smaller in comparison to the other sample types (e.g. up to 16.1% explained by the vineyard and up to 2.7% explained by the year).

Collectively, these results highlight the spatio-temporal variation in the vineyard microbial communities, with distinct patterns emerging for different sample types and microbial groups.

### The Fungal Berry Microbiome Is Strongly Shaped by Location and Shows a Robust Distance-Decay Relationship

Beta-diversity analysis revealed site effects to be the strongest driver of berry fungal community structure within AOC Lavaux across different years, despite a maximum inter-plot distance of only 2.46 kilometers. This spatial imprinting is strong enough (and temporally robust) that it enabled successful prediction of vineyard using Random Forest machine-learning classifiers trained on fungal community composition from one year and testing it on a different year. The area under the curve (AUC) values, ranging from 0.62 to 0.76 (see Supplementary Table 5), reflected the classifier’s ability to distinguish between sites, demonstrating moderate to strong predictive power in identifying the location based on the microbiome data between years. To further isolate the influence of this temporal variation we performed nested cross-validation (NCV) random forest classification using only samples from harvest 2021, achieving a mean AUC of 0.84 (Figure 2). Hence, microbial compositional similarity also appears to display temporal decay, with predictive accuracy of spatial location gradually decreasing from AUC of 0.84 within the same year (Figure 2) to 0.74 (2022) and 0.62 (2023) in successive years (Supplementary Table 5). The overall high predictive accuracy underscored the plot-specific signatures on the fungal community composition, while highlighting the relationship of vineyard location and vintage.

**Figure 2.**
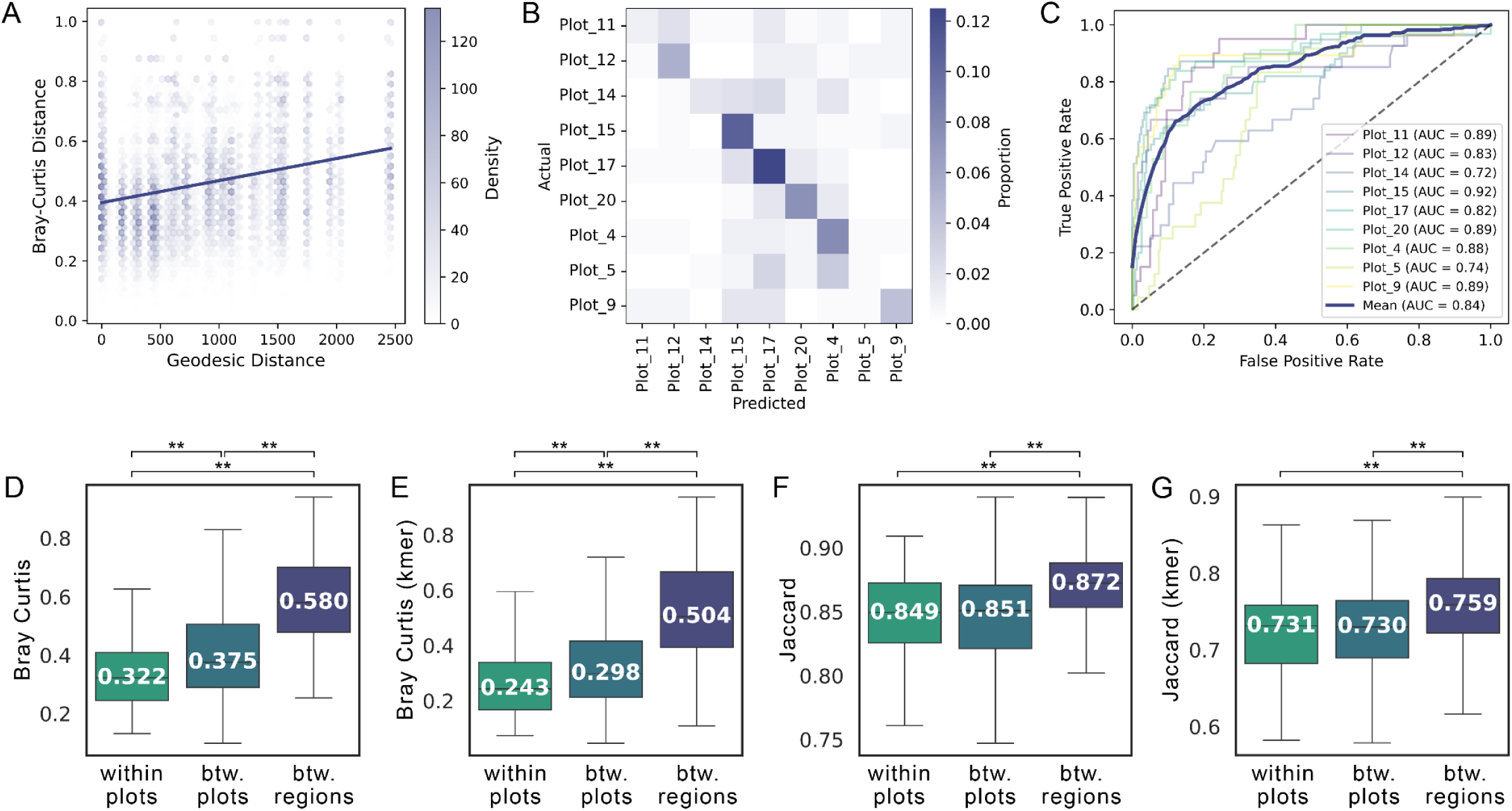
Berry fungal microbiome variation by site. Vineyards within the same region exhibit a strong distance decay relationship, as shown by a Mantel test using the geodesic distance (in meters) against Bray-Curtis distance between berry fungal communities from the 2021 harvest. (B-C) Prediction of site from fungal communities with a nested cross-validation random forest classifier (plots with fewer than 10 samples were excluded), as shown in the confusion matrix (B) and receiver operating characteristic (ROC) curves for the classifier performance (C). This is further reflected when comparing beta diversity distance within the same vineyard, or between vineyards of the same region or vineyards of different regions, using Bray Curtis (D), k-mer based Bray Curtis (E), Jaccard (F), and k-mer based Jaccard (G) metrics.

To further understand the underlying spatial heterogeneity, we analysed differences in fungal berry communities within vineyards as well as across regions. To capture intra-vineyard heterogeneity, we performed dense sampling in the AOC Lavaux vineyards at harvest in 2021, collecting samples approximately every 10th vine in every other row, excluding edge rows. To compare berry microbiomes across regions, in addition to samples from AOC Lavaux, we collected samples from the adjacent AOC Valais of Chasselas and Pinot Noir vineyards at harvest in 2023 (see Supplementary Figure 2). First, we compared various beta diversity metrics of the same grape variety from (i) within the same vineyards, (ii) to vineyards within the same region and (iii) vineyards from different regions (Figure 2). This showed that fungal communities from vineyards of different regions are robustly more dissimilar, than when comparing within the same region. This heterogeneity was more pronounced when taking the abundance of taxa into account. Using pseudo-phylogenetic metrics, i.e. considering the relatedness of present taxa, did not drastically change the results, indicating that the diversity between sites is not driven by differences in fungi belonging to entirely different clades but rather by closely related sub-species or strains.

Next, we evaluated the relationship between geographic proximity and community similarity. This revealed a robust distance-decay relationship in fungal berry microbiota, where spatially closer vineyards exhibited more similar fungal communities (Figure 2, Bray-Curtis distance; Supplementary Table 6, all beta diversity metrics). This trend was observed even when comparing across vintages, indicating a strong and consistent influence of location on community assembly, despite significant inter-annual variations (Supplementary Table 6). While broad-scale (inter-vineyard) spatial effects were substantial, fine-scale (intra-vineyard) variation was generally weak (Supplementary Table 7). Only two vineyards, Plots 14 and 17, exhibited significant internal distance-decay relationships. While these plots do not stand out in terms of their size or topographical features, the surrounding land use and human activity

might play a role in causing this internal spatial heterogeneity. Most of the sampled vineyards in AOC Lavaux are exclusively surrounded by other vineyards, but Plot 17 is flanked by a high traffic road as well as a railway. While we are not able to conclusively determine the underlying reasons, it is noteworthy that the berry fungal microbiome can exhibit significant distance-dependent gradients even at small scales.

Additionally, spatial distance remained a significant predictor of fungal community composition even when comparing across different vintages or grapevine varieties (Supplementary Tables 6 and 8). This robust spatial signal underscored the dominant influence of location on community assembly, which is consistent with our PERMANOVA results (Table 1). While host genotype (variety) alone did not significantly affect fungal community composition, in the context of the sample location it showed robust effects across all metrics (Table 3). This indicated that the host genotype effect is substantially smaller, and would be masked when not considering the sampling site.

**Table 3.**
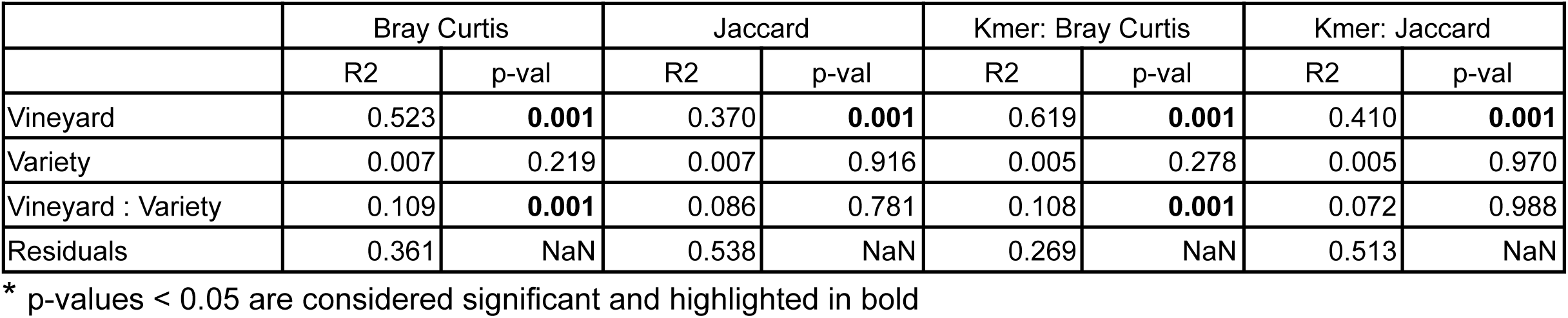
Permutational multivariate analysis of variance (PERMANOVA) with various beta diversity metrics for fungal communities of berry samples collected from AOC Lavaux as well as AOC Valais at harvest 2023.

|  | Bray Curtis |  | Jaccard |  | Kmer: Bray Curtis |  | Kmer: Jaccard |  |
| --- | --- | --- | --- | --- | --- | --- | --- | --- |
|  | R2 | p-val | R2 | p-val | R2 | p-val | R2 | p-val |
| Vineyard | 0.523 | **0.001** | 0.370 | **0.001** | 0.619 | **0.001** | 0.410 | **0.001** |
| Variety | 0.007 | 0.219 | 0.007 | 0.916 | 0.005 | 0.278 | 0.005 | 0.970 |
| Vineyard : Variety | 0.109 | **0.001** | 0.086 | 0.781 | 0.108 | **0.001** | 0.072 | 0.988 |
| Residuals | 0.361 | NaN | 0.538 | NaN | 0.269 | NaN | 0.513 | NaN |

\* p-values < 0.05 are considered significant and highlighted in bold

### Disentangling Environmental Factors Shaping Berry Microbiome Assembly: The Prominent Effects of Climate and Topography

With location having a dominant effect on fungal berry microbiome assembly, we further attempted to disentangle the influence of specific environmental factors. The vintage was the second most important factor in explaining variation in fungal communities (up to 2.7%, see Table 1), and we therefore analysed the climatic difference over the respective growing season for each year. Likely due to the close proximity of vineyards in Lavaux, we did not observe any significant differences in temperature or relative humidity when comparing within the same year (Supplementary Figure 3). With climatic differences being more pronounced between years, their effect on berry associated fungal communities is substantial. Relative humidity and temperature explained a variance of up to 4.2% and 0.6%, respectively (Supplementary table 9, Supplementary Figure 4). Yet, the specific climatic conditions of each plot, even subtle climatic variations, contributed to the observed biogeographic microbial patterns which becomes apparent considering the climatic influence nested per year, where relative humidity and temperature still significantly explain a variance of 2.7% and 0.6% respectively (Supplementary Table 10). Notably, differences in relative humidity had a stronger effect, and influenced the overall community structure of fungi, while the temperature solely had a significant effect on the presence or absence of fungi. Kmer-based metrics increased the explained variance, indicating that climatic conditions influence the community composition also on higher taxonomic levels. This is corroborated by the observation that fungal communities of vineyards with more similar climatic conditions also exhibit phylogenetically more similar fungal communities (Supplementary Table 11).

Since different climatic conditions appeared to result in markedly distinct fungal communities, we attempted to predict the relative humidity and temperature from the community composition using the machine-learning framework RITME (v1.0.4) [29] which systematically optimizes feature transformation and model selection in relation to the target variable. We evaluated four distinct supervised learning models and an extensive hyperparameter search for model optimization (see Supplementary Methods and Supplementary Figure 5). For predicting the temperature the best performing model was an elastic net linear regression, on ILR-transformed ASVs and data selection based on variance quantiles (above the 80th percentile), which explained 67% of the variance (Figure 3). The optimized prediction of the relative humidity is derived from a random forest model with features aggregated on the family level and selected with a variance threshold (≥8.42e-05) without transformation, which explained 71% of variance (Figure 3).

**Figure 3.**
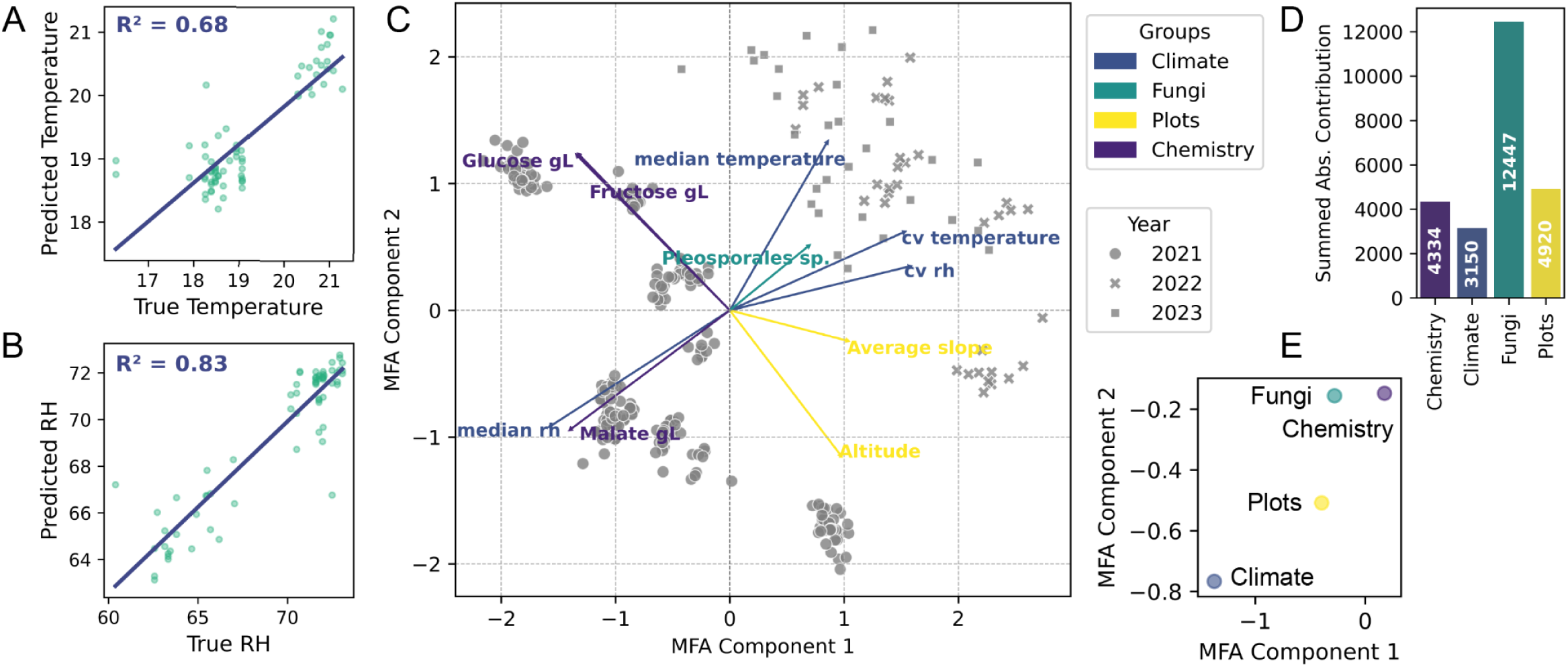
To predict the median temperature of the growing season we applied a linear regression model (A) and a random forest regression model to predict median relative humidity (RH). Multifactorial analysis (MFA) was used to disentangle the relationship between multiple data layers on the fungal microbiome of berries. Shown are biplots with the top 10 contributing features from all groups (C, colored by respective groups), as well the summed absolute contribution per group (D) and group representation (E).

To further study these complex interactions of key environmental and chemical factors influencing berry microbiome assembly, we conducted exploratory multifactorial analysis (MFA) integrating fungal microbiome data with plot topography, climate, and berry chemistry (sugars and organic acids). Despite the inherent complexity of the combined dataset, which explained only 18.56% of the variance with 10 components, the MFA revealed significant patterns. This low variance suggests that much of the variation remains unexplained, likely due to unmeasured factors, interactions, or the inherent heterogeneity of microbiome data. The MFA clearly separated samples by vintage, with 2021 samples clustering distinctly, likely due to lower temperatures and humidity compared to 2022 and 2023 (Figure 1, Supplementary Figure 3). This is supported by the MFA biplot, where temperature and humidity metrics (median and coefficient of variation) strongly contributed to the first dimension (Figure 3.A). The second dimension is driven by topographical features, highlighting how these factors collectively influence berry chemistry and fungal community structure, which are closely associated.(Figure 3E). While altitude and average slope were highly influential as individual factors, the combined contribution of all plot properties (including radiation and exposition) had the strongest overall influence on data structure (Figure 3B). In correlation analysis, we previously observed robust associations between many of these factors, for example altitude and sugar content, which in the MFA appeared as opposing drivers of the data structure. This model thereby helps reveal the interconnection of various environmental factors in ultimately conveying a highly vineyard specific imprint on fungal communities.

Overall, these results show that climate and topography collectively drive the dominant site-specific imprint on berry fungal communities and chemistry. Particularly relative humidity emerged as a key factor influencing fungal community structure, affecting the presence of microbes and composition even on higher taxonomic levels.

### Soil Microbial Communities Are Influenced by A Wide Array of Environmental Factors

The year, sampling time point, and location all were significantly associated with differences in the soil microbiome (Table 1 and Table 2). To better understand the influence of distinct environmental factors, we also performed MFA (Supplementary Figure 6). This analysis showed that, in contrast to berry communities, soil microbial communities exhibited less interannual variability caused by climatic factors, which is consistent with the PERMANOVA results (Table 2). The plot topography, specifically the slope, appears to be the most prominent contributing factor in shaping these vineyard soil microbiomes. Regarding the soil properties, the interconnected variables of clay content and soil waterlogging (hydromorphy), as well as soil organic matter content and carbon-nitrogen ratio are key factors influencing communities. The even distribution of variance among groups in the MFA suggests that these factors are collectively crucial for understanding the observed microbial patterns. Furthermore, distinct bacterial clades showed significant variation between plots, with notable differences observed in the families *Steroidobacteraceae*, *Vicinamibacteraceae*, *Gemmataceae*, and *Micropepsaceae*.

### Differential Abundance of Fungal Taxa and Fermentative Yeast Dynamics in Response to Environmental and Chemical Gradients

After demonstrating a robust association between environmental factors and berry mycobiome structure, we investigated the influence on specific taxonomic clades. Differential abundance analysis revealed associations between low and high relative humidity clusters with significant enrichment of *Sporidiobolaceae* and *Saccharomycodaceae*, while *Eurotiomycetes*, *Capnodiales*, and *Pleosporales* were depleted. Within *Saccharomycodaceae*, the fermentative yeast genus *Hanseniaspora* was specifically enriched (Figure 4).

**Figure 4.**
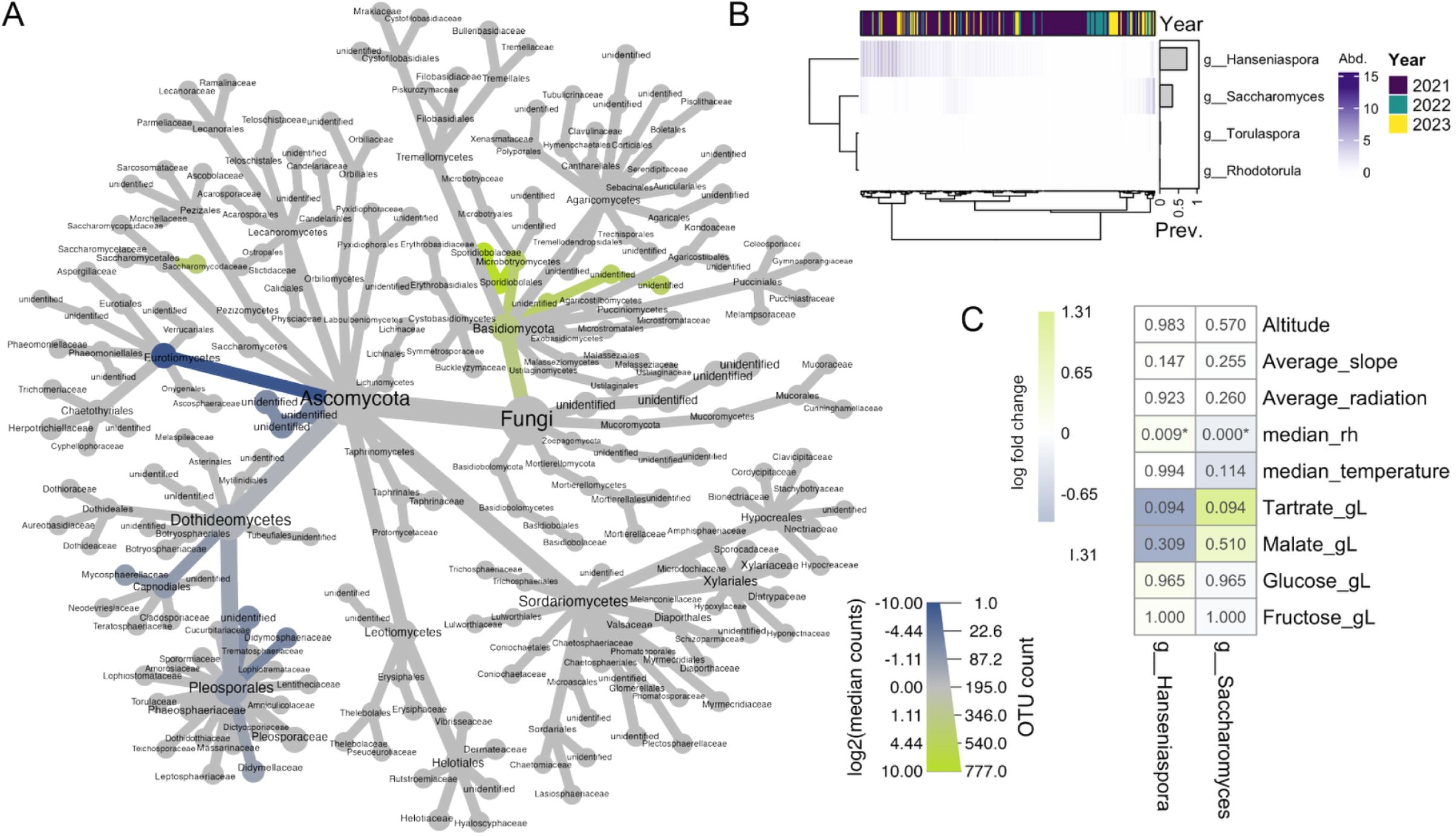
(A) The berry associated fungal communities displayed at the family level, exhibit significant differential abundance (enrichment in green, depletion in blue) between high and low relative humidity. Node size corresponds to overall feature count. (B) Log2-transformed abundance and overall prevalence of key fermentative fungal genera detected on berries at harvest across multiple years. (C) Differential abundance (ANCOM-BC2) analysis showing the log-fold change in bias-corrected abundance of Hanseniaspora spp. and Saccharomyces spp. associated with topographic, climatic, and berry chemical factors. Asterisks (*) denote statistically significant correlations (Holm corrected p-val < 0.05).

Among yeast species involved in subsequent wine fermentations, *Hanseniaspora uvarum* exhibited the highest prevalence, being detected in 0.72% of all ripe berry samples. It is followed by *Saccharomyces cerevisiae* (0.33% prevalence), whereas *Rhodotorula babjevae* and *Torulaspora quercuum* were sparsely detected (Figure 4). While most yeasts showed no significant inter-annual differences, *Hanseniaspora* spp. presence and abundance differed significantly between 2021 and 2022 (Fisher’s exact test for presence/absence, Bonferroni-corrected p = 0.0119; Wilcoxon Rank Sum test for log2-transformed data, Bonferroni-corrected p = 0.0117), suggesting that annual weather conditions could influence fermentative yeast prevalence and abundance in vineyards.

Next, we explored the influence of environmental factors and grape berry chemistry on the abundance of *Hanseniaspora* spp. and *Saccharomyces* spp. (Figure 4). Both yeasts exhibit a strong, yet non-significant, correlation with berry chemistry. Interestingly, this is an inverse relationship, whereby *Saccharomyces* spp. abundance was positively associated with organic acids and *Hanseniaspora* spp. with sugar content and relative humidity. This is consistent with the original observation of enrichment of *Hanseniaspora* spp. in vineyards with high relative humidity.

### Multi-Omics Analysis of Wines Highlights Vintage Effects and Plant-Microbe-Metabolite Interactions

After identifying distinct microbial clades, particularly fermentative organisms, that varied with environmental conditions, we next investigated the relationship between vineyard properties, environmental factors, and the berry microbiome in shaping wine characteristics. Therefore, we conducted microvinifications of grapes from each plot over three years. To replicate typical wine-making conditions and to ensure consistency across years, we inoculated the must with a white-wine-specific commercial *S. cerevisiae* strain. To preserve vineyard-associated bacterial communities, we did not add sulfur dioxide.

As expected, the microbiome during early fermentation is characterized by high diversity and abundance of vineyard-resident bacteria and fungi, but eventually alpha diversity declines as fermentative yeast and bacteria dominate the fermentations (Supplementary Figure 7). Beta diversity ordinations show a clear gradient in community composition throughout the fermentation (Supplementary Figure 7). Notably, when considering solely the presence/absence of taxa, the ripe berries and must already exhibited distinct microbial compositions, which is likely due to the introduction of winery-associated microbes.

To further explore the relationship between microbial communities and wine characteristics, we performed untargeted analysis of wine metabolites using three distinct methods: head-space gas chromatography-mass spectrometry (HS-GC-MS) for volatile compounds, and liquid chromatography-mass spectrometry (LC-MS) in both positive and negative ionization modes. The integration of these multi-omics data layers aims at providing a comprehensive framework for understanding the spatio-temporal influences on wine characteristics. Ordination analyses consistently revealed a clear clustering of wine samples by vintage across all data layers (Figure 5). This observed temporal stratification underscores the substantial impact of inter-annual variations on the microbial and metabolic composition of the wines, and reflects the dynamic nature of viticulture and winemaking. (Figure 5).

**Figure 5.**
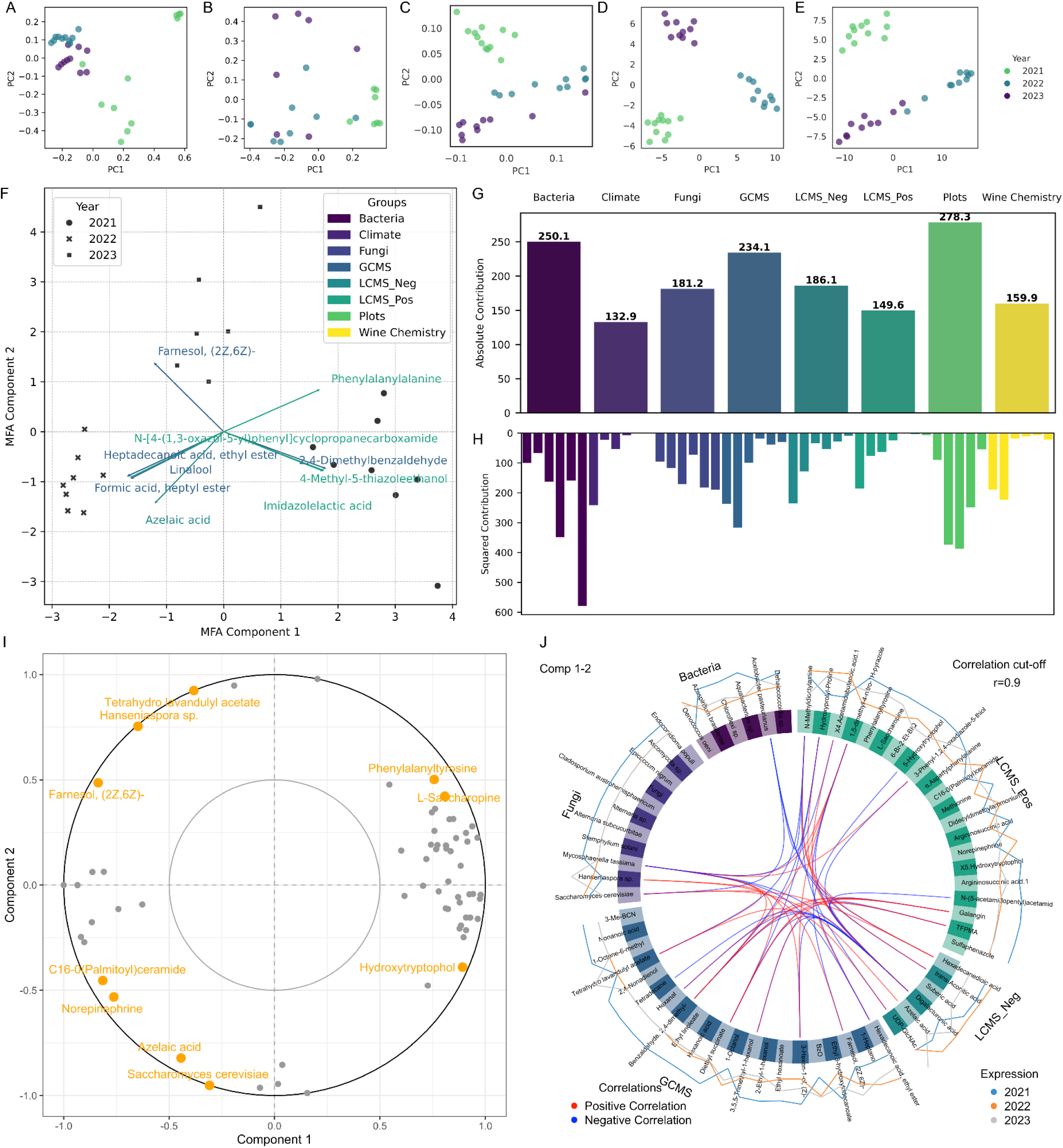
Ordination plots and multifactor analysis (MFA) to compare wine samples across different years with the various multi-omics datalayers. (A) PCoA plot for Jaccard metrics for fungal communities and (B) Jaccard metrics for bacteria, (C) PCA plot of head-space gas chromatography-mass spectrometry (HS-GC-MS) data, (D) PCA plot for negative mode liquid chromatography-mass spectrometry (LC-MS) and (E) PCA plot for positive mode LC-MS. (F–J) Exploratory MFA of wine samples with microbiome (fungal and bacteria),untargeted metabolomics (GC-MS, LC-MS), wine chemistry, climate data and topographical vineyard properties. (F) Shows the Biplot of top 10 contributing features in the first 2 dimensions as well as (G) the absolute summed contributions per group and (H) per dimension. Additionally, we performed a supervised, horizontal integration Partial Least Squares Discriminant Analysis (PLS-DA) model on the microbiome and metabolome data, (I) showing the variable plot with top 10 loadings driving the dispersion highlights in orange and (J) a circos plot of top correlations (cut off at 0.90).

To further study the interaction between multi-omics layers, vineyard topography, key wine chemistry parameters, and climatic factors, we conducted MFA using six components that explained 63.12% of the variance. The ordination plot of the first two components, accounting for 21.9% and 12% of the variation respectively, show a clear clustering by year, with various metabolites identified through GC-MS and LC-MS analysis driving the spread (Figure 5). While considering the absolute group contributions, the topographic properties of the vineyards had the greatest influence on the overall data structure. Notably, this contribution is most pronounced in the 3rd and 4th dimension of the model. In contrast, climate had the strongest influence on the first dimension, which may explain the horizontal spread observed between the samples from 2021 and 2022/2023, as these years had the largest climatic differences. These climatic variations are linked to corresponding differences in metabolites and wine chemistry, as these groups also are prominently associated with the first two dimensions of the model. Fungal and bacterial communities however show a more complex relationship to the other datalayers, as their group contributions are substantial across all dimensions. This suggests that these microbiomes are influenced by a combination of factors, including topography, the climate, as well as resulting wine chemistry and metabolites, which collectively shape their community structure.

While the unsupervised MFA revealed broad trends, we next applied a multiblock, supervised Partial Least Squares Discriminant Analysis (PLS-DA), specifically a Data Integration Analysis for Biomarker Discovery using a Latent Variable Approach for Omics studies (DIABLO), with two components. This approach allowed us to mine microbiome and metabolomics data for key discriminatory features and gain mechanistic insights. Using year as the target variable, we sought biomarkers that explain vintage-related differences. The variance between years was evenly distributed between the two model components (51.7% and 48.2%), confirming the robustness of the model in capturing group differences across the integrated datasets. PLS scores plots (Supplementary Figure 8) show strong clustering of each data layer by year, with more overlap in bacterial communities. The variable plot (Figure 5) highlights the most influential features driving the dispersion across components and their correlations. Most metabolites and microbes cluster to one side, with several key features standing out in driving the differences between years. Notably, we observed *Hanseniaspora sp.* and *S. cerevisiae*, each closely associated with various metabolites forming distinct clusters. This is also further illustrated in the circus plot, showing the top correlations and potential biological interactions among features (Supplementary Table 12).

Several intriguing correlations between the microbiome composition and wine metabolome indicate bi-directional interactions. A strong positive correlation was observed between *S. cerevisiae* and LC-MS-detected acids, such as the likely plant derived dicarboxylic hexadecanedioic acid and azelaic acid, which is known to have antibacterial properties. This is consistent with a strong negative correlation between *Acetobacter pasteurianus* and *S. cerevisiae*, suggesting that plant metabolites, which suppress bacteria, may indirectly promote yeast growth. *Hanseniaspora sp.* was strongly associated with volatile compounds like Farnesol and Tetrahydrolinalyl acetate (Tetrahydro lavandulyl acetate), which are both floral scents. Interestingly, the model also revealed strong positive and negative correlations of various metabolites, among them diethyl succinate which plays a role in the fruity aroma profile of wines, with the common grapevine endophytic fungus and opportunistic pathogen *Mycosphaerella tassiana*. We also observe multiple correlations among metabolites involved in the plant stress response, many of which are aromatic compounds, such as Hexanal, Farnesol, Lavandulyl-Tetrahydro, and Diethyl succinate (Supplementary Table 12).

In summary, by integrating environmental and multi-omics data, we highlight the complex interactions that shape the berry microbiome and metabolome. The analysis of specific interactions between bacteria, fungi, and metabolites sheds light on the plant’s biotic stress response, the interplay between microbes, and the direct link between fungal species and aroma formation in wine fermentation.

### Integrating Sensory Data Reveals the Complex Interplay of Factors in Shaping Wine Characteristics

Having shown the strong correlation between fermentative yeast species and volatile aromatic compounds, we further explored olfactory differences in wine characteristics through sensory analysis. A panel of experts evaluated each wine based on a set of predefined criteria typical of Chasselas wines from AOC Lavaux. Although there were substantial and significant differences between assessments (see Supplementary Figure 9), the wines from different vineyards still showed distinct flavor profiles (Figure 6). We integrated environmental factors, microbiome and metabolome data from the respective year as well as the sensory data using MFA, with a strong fit (four components explaining 77.19% of the variance). The contributions of the various groups were relatively balanced. Climate overall showed the smallest, yet most distinct effect on the first dimension. This aligns with previous observations that climatic differences between years were more significant than those within a single year within the AOC region. The biplot of top loadings further highlights the strong correlation between specific metabolites, most of which were measured in the LC-MS positive ionization mode.

**Figure 6.**
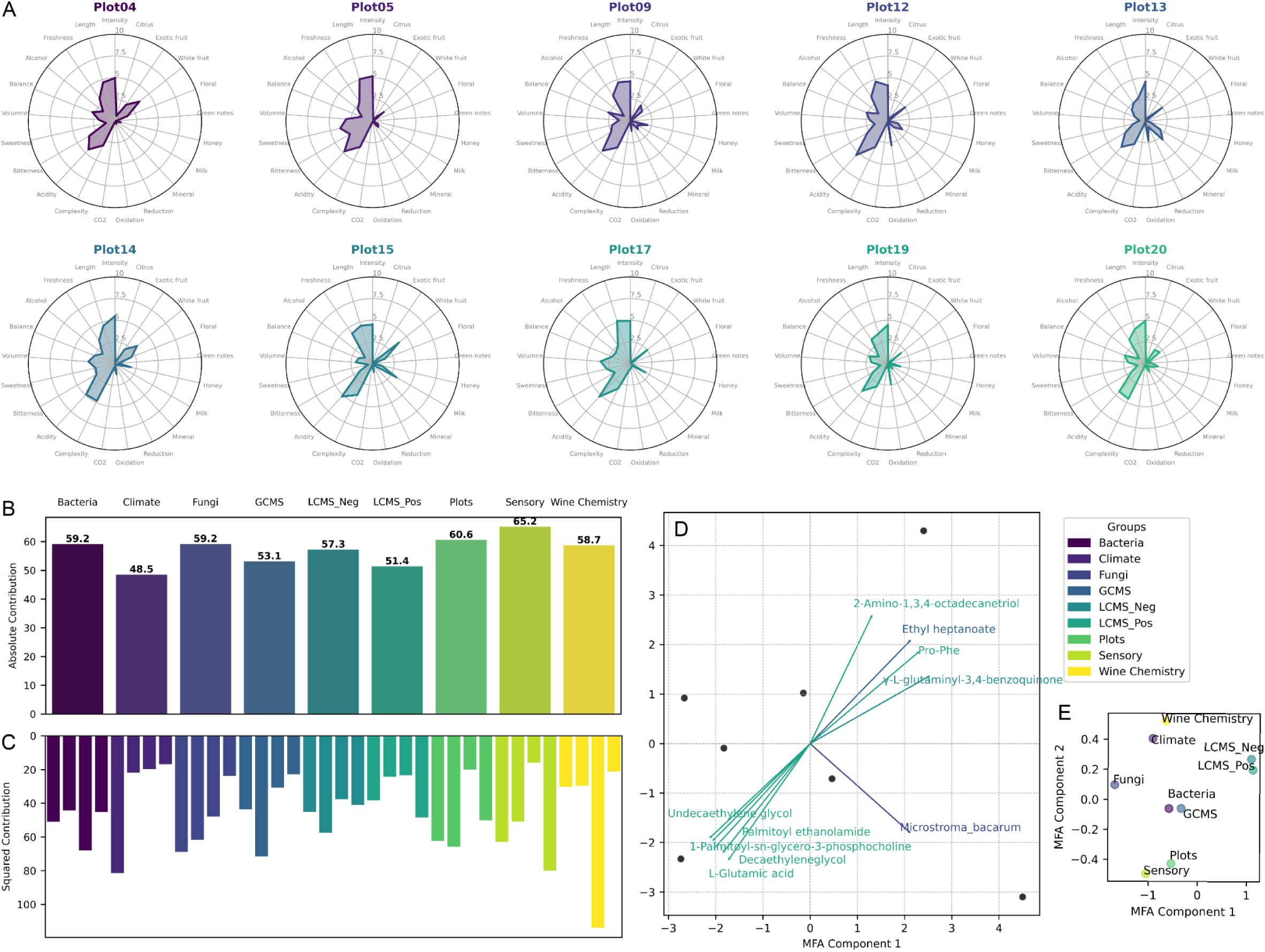
Integrative Analysis of Multi-omics and Sensory Data: (A) sensory spider plots illustrating the distinct characteristics of wines produced from different vineyards. (B) Multifactor analysis (MFA) results showing the summed contribution of each group to the model, as well as (C) contributions of each group per dimension. (D) The biplot with top 10 loadings reveals the association between specific metabolites and olfactory features, highlighting their role in driving the dispersion of wine samples. (E) Group representation shows the clustering of groups, highlighting their close interconnectedness.

Further investigating these correlations revealed strong associations between topography, climate and sensory properties. Specifically, higher altitude is strongly associated with lower median temperature (Spearman correlation strength = -0.96, p = 0.0005) and higher temperature variability (corr = 0.89, p = 0.0068), which in turn appear to influence sensory properties. Warmer and more stable climates enhance volume in the mouth (corr = 0.86, p = 0.0137), aromatic length (corr = 0.93, p = 0.0025), acidity (corr = 0.93, p = 0.0025), and sweetness (corr = 0.86, p = 0.0137), while greater temperature variability reduces these attributes (corr = -0.86 to -0.82, p < 0.025). Southwest-facing exposure was associated with reduced green notes (corr = -0.79, p = 0.034) and olfactory complexity (corr = -0.79, p = 0.034), likely due to increased sun exposure promoting ripeness. Notably, the two sensory characteristics, exotic fruit and balance, are not significantly associated with any climatic features, topography, or other fungal taxa, but exclusively with the fermentative yeast *Hanseniaspora spp.* (corr = -0.82 and -0.81, p-values = 0.023 and 0.027, respectively). *Mycosphaerella tassiana*, and other putative pathogenic fungi (*Alternaria spp, Stemphylium solani, Ustilago hordeii, Cryptovalsa ampelina, Endoconidioma populi, Curvularia trifolii*) were shown to strongly positively correlate with the sensory perception of Oxidation (corr = 0.75 to 0.91, p < 0.05), but commonly had an equally strong negative correlation with Olfactory Intensity (corr = -0.75 to -0.81, p < 0.05) in resulting wines.

## Discussion

The concept of *microbial terroir* describes the influence of site-specific microbial communities on distinct product characteristics and has been reported for a variety of foods [30–32], but plays a particularly prominent role in viticulture [5,33–35]. Microbial communities in vineyards influence plant health [36] and berry qualities, as well as contribute directly in subsequent fermentation [4,14]. Despite the location commonly being the primary differentiating factor between vineyard microbiomes [1], the role and interaction of specific underlying environmental drivers in shaping associated communities, and the direct impact on resulting wine characteristics remain poorly understood.

Our study provides a more granular and comprehensive perspective by disentangling location-dependent environmental factors within a tightly controlled and spatially linked experimental framework that controls for biotic factors. This allows us to dissect associations between specific environmental factors and grapevine- and wine-associated microbiota and metabolomes. Even within a small geographic region, we observed highly distinct microbial communities, shaped by site characteristics, annual and seasonal fluctuations, and microclimatic conditions.

Soil is a fundamental element in *terroir*, providing the vine with water and nutrients, and in the context of *microbial terroir*, acting as a primary microbial reservoir. It serves as a key source for the colonization of annually growing leaves and berries [12,37], with microbial transmission occurring through various potential vectors [1]. In our study, soil microbial communities exhibited strong spatial structuring, forming the foundation for subsequent heterogeneity in other vineyard-associated microbiomes. At the intra-region scale (max 2.46 km distance between vineyards) soil microbiome variation was not driven by geographic proximity, as was reported in studies across larger distances [38]. This is particularly interesting, as in berries we observe a strong distance decay relationship, indicating distinct transmission, dispersion and environmental filtering mechanisms for these different microbial habitats. Microbial diversity in soil was driven by intricate interactions between inherent soil properties, such as clay content and connected water logging, as well as dynamic factors like soil carbon and nitrogen levels. These factors all contributed to observed seasonal fluctuations, which was also reflected in soil pH - a key variable which is influenced by climate, altering microbial activity and thereby the organic matter content [39]. In contrast, microbes colonising aboveground plant organs are exposed to harsh conditions, such as UV radiation and fluctuations in temperature and humidity [1], imposing strong selection pressures and shaping the composition of epiphytic communities.

As such, and most similar in richness and community composition to soil, bark is also a key microbial reservoir. Fungal communities of bark showed significant spatial differences, which were stable across the years. This may be connected to long-term niche adaptation, as fungi form stable communities in the rough surface of bark, which become more complex as the plant ages [40], as well as the dispersal limitation of fungi leading to more spatial heterogeneity [33]. Conversely, bacterial communities on bark varied more strongly between years than between sites, indicating a more dynamic response to climatic fluctuations.

Fungal communities on leaves and berries, as annually newly formed and colonized organs, are as expected variable between the years. While before ripening, the berry microbiome closely resembles that of leaves [41], the accumulation of sugars and other metabolites leads to a niche specialization in later stages [1]. Interestingly, while leaf microbiomes were not significantly different between vineyards, the berry-associated fungal communities exhibited the strongest spatio-temporal partitioning of all sample types. The site specific imprint was notably strong, with some vineyards even exhibiting significant internal heterogeneity. Predictions of the vineyard location from the fungal berry microbiome improved when controlling for host genotype or vintage, or comparing across larger geographic scales, which highlights the mutual influence of host genotype and site-specific factors on community-assembly. The site-specific factors driving this diversity of berry fungal microbiomes are primarily attributed to climatic and topographic variation between vineyards. Larger inter-annual climatic differences had a corresponding larger influence, yet even insignificant microclimatic differences between plots showed to have a significant effect in structuring fungal communities, with climatically more similar plots also harbouring more similar fungal communities. Particularly relative humidity had a strong effect on taxonomic community structure, including a noteworthy association with fermentative organisms *Hansensiaspora spp.* and *Saccharomyces cerevisiae*. Notably this appears to be an inverse relationship, with climatic factors and resulting berry chemistry either favouring *Hansensiaspora spp.* or *Saccharomyces cerevisiae*, which also showed to be a key distinction between wines of different years.

These observed microbiome variations between plots and years were further reflected in distinct metabolomics profiles of the resulting wines. Inter-annual climatic variations as well as fine-scale site-specific differences contributed to distinct characteristics. We found that *Hanseniaspora spp.* are strongly associated with the expression of floral aroma compounds across vintages, and distinctly correlated with exotic fruit aroma and balance of samples from the same year. Previous lab-controlled studies [42,43] have shown that fermentation with *Hanseniaspora* species contributes to more fruity and floral aromas, with species and strain variability playing a significant role in the formation of specific aroma compounds [44]. *Hanseniaspora* spp. have therefore been suggested to be major contributors to *terroir* expression. These yeasts are particularly influential during the early stages of fermentation, as they are more alcohol-sensitive, underscoring the importance of fermentation practices on their aroma-producing potential [35]. Interestingly, the prevalence of *Hanseniaspora* species has been shown to vary between grape varieties [45], regions [35,46], vineyards [2], years [47] and with berry ripening and health status [41]. We contribute to this, by providing a promising, direct link between site-specific climatic factors, microbial differences, and fermentation metabolites, leading to sensory variations in resulting wine.

Notably, when comparing fermentation dynamics across different years, we observed a significant interaction between plant stress response metabolites, microbial communities and aroma formation. Specifically, higher levels of azelaic acid, a key signaling molecule involved in the expression of antimicrobial genes as part of the plant’s pathogen stress response [48], were associated with increased growth of *Saccharomyces cerevisiae*. Azelaic acid serves as a pathogen response signal and plays a role in lipid signaling related to grapevine defense against downy mildew [48] and esca disease [49]. Such associations between grape volatiles, lipid metabolism and the associated microbiome have previously been suggested [50]. Similarly, we observed multiple correlations among metabolites involved in plant stress response, many of which are aromatic compounds. Microbes, including yeasts and potential pathogens, have been shown to modify plant volatile compounds in grapevines [51] and our findings highlight these interactions between microbes and plant biotic defense mechanisms contributing to wine aroma formation.

Additionally, our results highlight an important but often overlooked aspect of *microbial terroir*: the impact of vintage. While it is well understood in winemaking that vintage influences wine quality, there is a common misconception that microbial communities should remain stable if they are truly regional. In reality, these communities are shaped not only by location but also by annually variable environmental conditions. As we show, 2021 differed markedly from 2022 and 2023, which were climatically more similar. Vineyard microbiomes – and the resulting wine characteristics – respond to these fluctuations and we observed a form of temporal decay, where microbiomes from consecutive vintages tend to be more alike. It is therefore essential that future studies consider not only spatial but also temporal variation, recognizing *microbial terroir* as a dynamic expression of place and time.

Particularly when studying plant-microbe interactions or fermentations, strain diversity plays a crucial role, and within-species variation can influence whether a microbe contributes to aroma formation or spoilage, or acts beneficial or pathogenic [52]. Our analysis was conducted primarily on the level of amplicon sequencing variants. While the large number of samples provides robust insights, we lack the resolution to track the transmission of specific strains between plant organs, fermentation stages, and across time and space. Another strength, yet in this context a limitation, of our study is the setting in commercially managed vineyards. While this conveyed insights into microbial diversity in real-life settings, we could not control the application of fungicides and herbicides as well as under-vine management practices. Showing the connection between environmental factors, plant stress, associated microbes and resulting wine characteristics therefore calls for future studies systematically exploring the interactions with vineyard management, in particular in relation to abiotic plant stress.

Overall, our study provides insight into complex microbial dynamics in vineyards and subsequent fermentation, and successfully disentangles abiotic contributions in highly interconnected systems. We advance the understanding of *microbial terroir* in viticulture and winemaking by showing how fine-scale environmental variation shapes microbial diversity and drives interactions between the microbiome and metabolome of grapevines and wine.

However, establishing causal relationships among these factors remains a key challenge. The *microbial terroir* research community must also recognize the need to control many covariates for optimal study design, and adresse the challenge of controlling these factors in commercial vineyards. Future research will require higher-resolution sequencing technologies, larger sample sizes, and higher temporal resolution to track transmission of strains and metabolomics profiling to fully reveal these intricate interactions.

## Supporting information

Supplementary Methodology

## Supplementary Material

- Supplementary Methodology
- Supplementary Figures and Tables

## Data Availability

Amplicon sequencing data is available from the Sequence Read Archive (SRA) under accession number PRJEB89111 (16S) and PRJEB89112 (ITS).

## Code Availability

All code used in this study is available on GitHub:https://github.com/LenaFloerl/microterroir.git.

## Acknowledgements

We would like to thank the winegrowers of the Villette AOC Lavaux and Valais for letting us collect samples over 3 years. Further we thank Vivian Zufferey and Jean-Sébastien Reynard of Agroscope for collecting samples in Valais, Serafina Plüss and Julinka Wäsche for technical support, Ondino Azario and Laura Nyström (ETH Zürich) for supporting the GC-MS analysis, and Alfonso Die for supporting HPLC analysis.

The authors thank the Genetic Diversity Centre (GDC) of ETH Zurich for supporting the library preparation. The microbiome amplicon sequencing and LC-MS metabolomics analysis was performed at the Functional Genomics Center Zurich (FGCZ) of University of Zurich and ETH Zurich.

The authors gratefully acknowledge financial support from the Swiss National Science Foundation [Grant Number: 310030_204275] (to NAB) and the Swiss Government Excellence Ph.D. Scholarship (to LF).

## Author Contributions

The study was conceived by NAB and MR, and supervised by NAB. LF and PS collected samples, and PS recorded viticultural metrics and performed the microvinifications. LF processed the samples, performed the analysis, and wrote the article with review and contributions from NAB, MR and PS.

## Abbreviations

ANOVA: Analysis of Variance
AOC: Appellation d’Origine Contrôlée
AUC: Area under the curve
cv: coefficient of variation
DIABLO: Data Integration Analysis for Biomarker Discovery using Latent Variable Approach for Omics studies
FTIR: Fourier Transform Infrared Spectroscopy
HPLC: High pressure liquid chromatography
HS-SPME-GC-MS: Headspace solid-phase microextraction gas chromatography–mass spectrometry
LC-MS: liquid chromatography–mass spectrometry
MFA: Multifactor analysis
PERMANOVA: Permutational multivariate analysis of variance
RH: Relative humidity
RT: retention time
TIC: total ion current
VOCs: volatile organic compounds
YAN: yeast-assimilable nitrogen

