## Supplementary Methodology for "Grape Expectations: Disentangling Environmental Drivers of Microbiome Establishment in Winegrowing Ecosystems"

#### 1. Longitudinal Sampling and Environmental Data Collection in AOC Lavaux

In the AOC Lavaux subregion Villette, we collected samples and data over 3 years (2021 to 2023) from 12 different vineyards, two of which had their vines uprooted within this time span. The vineyards are all within a 2.5 km radius and were selected to represent maximal pedoclimatic variability present in the AOC region. They are uniformly planted with *Vitis vinifera* clonal variety Chasselas on a 3309C rootstock. Vine age ranges from 15 to 25 years and all vines are guyot-pruned and trained in a vertical shoot positioning system [1]. Over the growing seasons we recorded vineyard management and vine development, i.e. application of herbicide, hail damage, mildew appearance, estimated grape yield and pruning weight, as well as records of phenological stages of budburst (BBCH 09) and flowering (BBCH 65) on 100 randomly selected plants within a plot [2].

To record climate data we installed sensors (HOBO U23 Pro v2 Temperature/Relative Humidity Data Logger from Onset, Cat No. U23-001A) with radiation shields (Onset, Cat No. RS3-B) for improved temperature measurements at approximately 1.8 m height within each vineyard plot. These continuously measure the temperature and relative humidity every hour for 3 years. We removed outliers of the climate data (generated by e.g. dropped probes, empty batteries) with rolling Median Absolute Deviation (MAD) and subsequently calculated the daily median values over each growing season (1. April to 30. September). Additionally we calculated the Growing Degree Days (GDD) for each plot and year.

Samples for microbiome analysis were collected over a three-year period. Rhizosphere samples were taken twice per year (in spring and at veraison), while bark and leaf samples were collected annually at veraison. Berry samples were collected twice per year, at veraison and harvest. From the Valais vineyards we collected berries in 2023 at harvest. To collect rhizosphere samples, the surface layer of soil (app. 3 cm) was removed and three soil cores were then taken in proximity to the stem (within 20 cm from the trunk) to 30 cm depth. Soil samples were pooled and sieved with a 1 mm mesh size to remove stones. For grape sampling, clusters of berries were cut off using sterile scissors. Bark and leaf samples were also taken using sterile scissors, ensuring no contamination occurred during the sampling process. All collected samples were transported on ice back to the laboratory and stored at −80 °C until further processing.

### 2. Berry Chemistry

To assess berry maturity we used High pressure liquid chromatography (HPLC) to analyse the concentrations of glucose, fructose, malic- and tartaric acid in grape must samples [3]. Glucose and fructose are essential for fermentation of grape juice to wine, and are an indicator of berry ripeness. Tartrate and malate were investigated because they are the most prominent organic acids in wine making up 90 % of total acidity in grape juice [4].Therefore we used diluted 500 µL diluted (1:5, with ultrapure Milli-Q water) and and filtered (0.2 µm) grape must in HPLC closed glass vials (e.g. crimp neck from BGB Analytik AG, Cat No. 0800035 and alu crimp caps with septum from BGB Analytik AG, Cat. No. 080304) and put them in an autosampler (Merck Hitachi, No. L-7250) . To calculate the concentration of each component a calibration curve with solutions of the named sugars and organic acids was used. All standards, e.g. D-(+)-Glucose 99.9% (Cat No. PHR1000), D-(−)-Fructose ≥99% (Cat No. F0127), DL-Malic acid ≥99% (Cat No. 240176), L-(+)-Tartaric acid ≥99.5% (Cat No. T109), were purchased from Sigma Aldrich. Samples were run on a HPLC-UV-RI Agilent 1260 Infinity with a carbohydrate column HPLC Aminex HPX-87H (300 x 7.8 mm) column (Biorad, Cat. No. 1250140) and a guard column e.g. Cartridges Carbo-H (4 x 3.0 mm) (Phenomenex, Cat No. 125-0129). The column is heated to 35°C (column oven from Merck Hitachi, No. L-7300), 20 µL of the samples are injected into the system and eluted with 0.65 mM H2SO4 at a flow rate of 0.5 mL/min (pump from Merck Hitachi, No. L-7100). UV detection is carried out at 210 nm and analysis in the EZChrom Elite software (Agilent).

Further we measured the Brix value of berries using a handheld refractometer for sugar (VWR Cat. No. 635-0631, 0-32 % Brix, 0,2 °Brix resolution). To assess vine water status and photosynthetic carbon isotope composition as a proxy for vine water stress, the 12C/13C ratio (δ13C) in sugars of ripe berries was analyzed in 2023 by Laboratories Dubernet in Narbonne, France. Additionally, yeast-assimilable nitrogen (YAN) was measured by Fourier Transform Infrared Spectroscopy (FTIR) at Changins.

### 3. Soil analysis

To measure the pH of each soil sample we followed the Standard Operating Procedure by the FAO  [5]. In short, we used 0.35 g of soil with 1.75 µL of 0.01 M CaCl2 (Sigma Aldrich, Cat No. 1.02382) solution in a Deep Well plate, which is homogenised for 30 minutes at 8 Hz in a TissueLyser (QIAGEN) before let settle for another 30 minutes and measured with a calibrated pH meter.

The Sol-Conseil laboratory in Changins analysed soil samples collected from 10 plots in 2023. Samples were conditioned (dried for 48h at 40°C and sieved by 2mm) and subsequently analysed for total nitrogen content, and C:N ratio as well as particle size distribution [Sand/Silt/Clay], including total organic matter (granulometrie).

#### 4. Sample processing and DNA extraction

For DNA extraction, the grapes were thawed and pressed using a stomacher. A 250 µL portion of the homogenized grape material was used for DNA extraction using the MagAttract PowerSoil Pro DNA Kit (QIAGEN, cat. no. 47109) and the PowerBead Pro Plate (QIAGEN, cat. no. 19311), following the manufacturer's instructions. Cell lysis was performed using a Bead Mill (Retsch), and DNA extraction was carried out on a KingFisher Flex (Thermo Fisher Scientific). The extracted DNA was stored at −20 °C until further processing.

#### 5. Library Preparation and Sequencing

Using the HighALPS ultra-high throughput library preparation protocol [6] which based on a unique dual index (UDI) strategy with custom 12 nt long Golay barcodes, we profiled present bacterial communities with amplicon sequencing of the hypervariable V4 region of the 16S rRNA gene using the updated 515F primer (5'-GTGYCAGCMGCCGCGGTAA-3') [7] and 806R primer (5'-GGACTACNVGGGTWTCTAAT-3') [8]. For fungal communities we amplified the first loci in the internal transcribed spacer (ITS) region with above described BITS and B58S3 primers [9]. To amplify the low microbial biomass samples we performed a nested PCR. In the first PCR all template DNA is enriched with 515F/806R primers and in the second PCR the amplicons are UDI barcoded as well as plant host plastid and mitochondrial 16S rRNA gene contaminations depleted with peptide nucleic acid (PNA) PCR clamps [10,11]. Therefore we first set up a 25 µL PCR reaction with 12.5 µL of 2x KAPA HiFi HotStart ReadyMix (Roche, Cat. No. 07958935001), 0.4 µM of each primer (Microsynth) and 2.5 µL of extracted DNA. All PCR reactions were set up using an epMotion liquid handling platform. Reaction conditions were initially 95 °C for 5 min, followed by 35 cycles of 95 °C for 20 sec, 49 °C (ITS) / 55 °C (16S) for 15 sec, and 72 °C for 15 sec, and a final extension of 72°C for 5 min. From this enriched template we used 1 µL per 25 µL reaction for a second PCR with 12.5 µL of 2x KAPA HiFi HotStart ReadyMix, 0.4 µM of UDI primer combinations as well as 0.5 µM or mPNA and pPNA clamps. The PNAs were vortexed and incubated at 60 °C for 10 min before use. The subsequent reaction conditions were 95 °C for 5 min, followed by 8 cycles (ITS) and 12 cycles (16S) of 95 °C for 30 sec, 78 °C for 5 sec for PNA annealing (only for 16S), 49 °C (ITS) / 50 °C

(16S) for 20 sec, and 72 °C for 15 sec, and a final extension of 72°C for 5 min. Notably, the presence of PNAs in the reaction showed that a lower annealing temperature increased yield. Resulting DNA concentrations were measured in duplicates with Qubit dsDNA High Sensitivity Assay (Thermo Fisher Scientific, Cat No. Q32854) on a Tecan Spark Microplate Reader. Amplicons were pooled equimolarity in two separate pools with a liquid handling platform (Brand GmbH, Wertheim, Germany) and purified with Agencourt AMPure XP magnetic beads (Beckman, Cat No. A63882) with a ratio of 0.7 X to remove primer dimers and small fragments on the KingFisher Apex. Quality of the resulting amplicons was controlled on a Tapestation (Agilent). To increase yield we reconditioned each pool separately with 12.5 µL of 2x KAPA HiFi HotStart ReadyMix, 0.4 µM of the standard Illumina P5 (5′-AATGATACGGCGACCACCGAGATCT-3′) and P7 (5′-CAAGCAGAAGACGGCATACGAGAT-3′) PCR primers and 2 µL template of the bacterial pool and 3 µL template of the fungal pool. Reconditioning PCR conditions were set to initial 95 °C for 3 min, followed by 4 cycles of 98 °C for 20 sec, 62 °C for 15 sec, and 72 °C for 30 sec, and a final extension of 72°C for 1 min. Ultimately we conducted a two-sided clean up with 0.2 X magnetic bead ratio, followed by 0.7 X magnetic bead ratio of the supernatant to remove remaining genomic DNA as well as primers from the reconditioning. The pools were combined and submitted to the Functional Genomics Center Zürich for Paired End 250 bp sequencing on the Illumina NextSeq 2000 P1 600 cycles with 20 % PhiX.

**6. Microbiome Data Analysis**

Microbial diversity analysis was performed with QIIME 2 (version 2024.10) [12]. Demultiplexed Illumina FASTQ files were imported, and residual adapter sequences were removed with cutadapt [13]. Amplicon sequence variants (ASVs) were generated by denoising single-end reads with DADA2 [14]. Taxonomic classification of ASVs was performed with the q2-feature-classifier plugin [15]. For 16S rRNA gene ASVs, a naive Bayes classifier was used, which was trained against the 99% SILVA 138 release database [16], curated with RESCRIPt [17], trimmed to the V4 region (515F-806R) and weighted [18] to improve representation of plant-surface communities. Fungal ASVs were classified using the UNITE database (version 9.0, released 2023-07-18) [19]. Contaminant ASVs were identified and removed using decontam, based on DNA extraction negative controls. Additionally, non-target ASVs were filtered: specifically, fruiting body-forming mushrooms were removed from the ITS dataset, and host DNA sequences (mitochondria, chloroplasts) were removed from the 16S dataset. All ASVs that could not be classified to at least the phylum level were also extcluded. For downstream diversity analyses, samples were rarefied separately by sample type and amplicon, with rarefaction depths specified in Methodology Table 1. Non-phylogenetic alpha (Pielou evenness [20], observed features, Shannon entropy [21]) and beta diversity metrics (Bray Curtis [22], Jaccard [23]) were calculated using the q2-diversity plugin. Additionally, k-mer based diversity metrics were calculated [24]. For 16S rRNA gene data, a phylogenetic tree was constructed using fasttree [25] and MAFFT alignment [26] with the q2-phylogeny plugin, and this tree was used to calculate phylogeny-informed diversity metrics (weighted UniFrac, unweighted UniFrac [27]) with q2-diversity.

*Methodology Table 1. Rarefaction depths for various sample types and amplicons and the thereby retained features and samples.*

| Sample Type | Depth | Retained |
|---|---|---|
| **ITS** | | |
| Bark | 15’000 | 450’000 (45.13%) features in 30 (90.91%) samples |
| Soil | 5’000 | 220’000 (30.45%) features in 44 (80.00%) samples |
| Leaves | 8’000 | 224’000 (33.53%) features in 28 (90.32%) samples |
| Berries | 10'000 | 3’400’000 (26.33%) features in 340 (95.51%) samples |
| Microvinification | 4'500 | 423’000 (22.68%) features in 94 (95.92%) samples |
| **16S** | | |
| Bark | 300 | 7’200 (0.72%) features in 24 (72.73%) samples |
| Soil | 1'500 | 70’500 (2.20%) features in 47 (85.45%) samples |

Permutational multivariate analysis of variance (PERMANOVA) was conducted using the ADONIS function [28,29] via q2-diversity. Mantel tests were performed using the vegan package [29], based on Spearman correlation with 1000 permutations.

For the prediction of the vineyard from fungal communities of berries we used a supervised Random Forest machine-learning classifier of the q2-sample-classifier plugin [15]. The models were trained with 10,000 trees. For training the classifier on one year to predict another we excluded plots that were not sampled across all three years (Plot 11, Plot 18). For the nested cross-validation assessing solely 2021 we excluded plots with fewer than 10 samples.

To predict climate variables from fungal berry microbiomes we used the machine-learning framework RITME (v1.0.4) [29]. We tested multiple transformation approaches to account for compositional data characteristics – including no transformation, presence/absence encoding, and log-ratio transformations such as Isometric Log-Ratio (ILR), Centered Log-Ratio (CLR), and Additive Log-Ratio (ALR) – as well as exploring different levels of taxonomic aggregation, ranging from ASVs to genus, family, order, and class. Additionally, RITME applies various filtering strategies based on variance and abundance, including thresholding, quantile-based selection, and top-ranked feature selection. We evaluated four distinct supervised machine learning models and types – linear regression, extreme gradient boosting, neural network regression, and random forest regression – using an extensive hyperparameter search for model optimization, and conducting 900 trials per model type for all berry harvest samples.

For microbiome analysis involving taxonomic assignment, we clustered ASVs against the respective reference database at 90% similarity. Multifactor analysis (MFA) was performed using prince [30]. Partial least squares discriminant analysis (PLS-DA) was conducted using the DIABLO function from mixOmics [31]. The DIABLO model showed high classification accuracy for LC-MS metabolomics data (positive ionization: 94.7%, negative ionization: 92.1%) and moderate accuracy for GC-MS (85.8%) and fungi (76.8%). Bacteria, however, showed lower classification accuracy (59.9%), indicating a weaker contribution of bacterial communities to the model. This is also reflected in the proportion of explained variance, where metabolomics and fungi contributed most to the model variance (Fungi: 52.4%, positive LC-MS 56.0%, negative LC-MC 40.7%, GC-MS: 46.7%), while bacteria accounting for the lowest variance (31.6%). HeatTrees were generated using Metacoder [32]. Additionally, differential abundance was analysed with ANCOM-BC2 [33]. All visualizations were generated using matplotlib [34] and seaborn [35]. Geographic maps were created using geopandas [36].

**7. Microvinifications**

Microvinifications were performed with five kilograms of Chasselas grapes (*vitis vinifera cv. Chasselas*) frome every plot, which were randomly collected a few days prior to commercial harvest. Grapes were kept at 4°C overnight. The next day, without destemming the fruit, the grapes were pressed twice for 1 min. using a stainless steel BagMixer® 400 (Interscience, St. Nom, France), the pomace was removed. A first series of samples was taken from the musts of each batch for analysis. At this point, no enological treatment was applied, and sulfur dioxide was not added to the musts to preserve the original microbiome from grapes to musts. The musts were then again kept at 4°C overnight for sedimentation before removing part of the sediments by a rough racking. Each must sample was filled into two magnum-sized bottles and immediately inoculated with the commercial *Saccharomyces cerevisiae* C19 yeast strain (Levuline C19, Oenofrance, France) which is commonly used in white wines to foster expression of terroir, to start the alcoholic fermentation. Fermentation temperatures were maintained at 18°C by adjusting the room temperature to that temperature. Must densities were monitored daily. After 14 to 18 days, as soon as alcoholic fermentations in the individual batches were completed, samples were collected for analysis and the fermented musts of each plot filled into three standard sized bottles and kept at 18°C room temperature to naturally induce malolactic fermentations. After several weeks, at the end of malolactic fermentations, a third series of samples was taken from each batch. A final sedimentation was carried out and sulfur dioxide (30 mg/L) was added before bottling the young wines. After a short storage period, a sensory evaluation panel evaluated and scored the wines.

**8. Sensory Analysis**

The sensory methodology employed was a descriptive test using a sensory profiling approach known as Quantitative Descriptive Analysis (QDA). The sensory profiling method allows quantification of the perceived intensity of several sensory attributes for each wine. A predefined list of sensory attributes typical for Chasselas from AOC Lavaux was established beforehand (Olfactory intensity, Citrus, Exotic fruit, White fruit, Floral, Green notes, Honey, Milk, Mineral, Reduction, Oxidation, Olfactory complexity, CO2, Acidity, Bitterness, Sweetness, Volume in the mouth, Balance, Alcohol, Freshness on the palate, Aromatic length). The wines were served in anonymous black INAO glasses (40 ml) identified by a three-digit code. They were served at a temperature of 12°C ± 1°C. A comparative profiling method was applied, where tasters had the opportunity to compare the wines against each other. They were required to evaluate the intensity of each sensory attribute on a 10 cm linear scale ranging from 'absent' on the left to 'very intense' on the right. Tasters marked the perceived intensity level for each wine. Scores were then transformed into ratings ranging from 0 (left endpoint) to 10 (right endpoint). The taster panel consisted of 15 trained panelists. Each taster evaluated the wines in sensory analysis booths compliant with current AFNOR standards. The results were analyzed using a 2-factor Analysis of Variance (ANOVA) (wine factor and judge factor) to identify differences between the wines for each sensory attribute.

**9. Untargeted Metabolomics with HS-SPME-GC-MS**

We used untargeted Headspace solid-phase microextraction gas chromatography–mass spectrometry (HS-SPME-GC-MS) to analyse volatile organic compounds (VOCs) in grape must and microvinification samples. The headspace analyses were conducted on a Trace 1310 GC (Thermo Scientific) gas chromatograph (GC) coupled to a TSQ 8000 EVO (Thermo Scientific) mass spectrometer (MS) equipped with an RSH (Thermo Scientific) autosampler. Each GC glass vial contained 1 mL of sample with 150 µL buffer (0.5 M $Na_2HPO_4*2H_2O$ , 0.1 M $KH_2PO_4$) and 0.5 g NaCl (>99.5 % purity, Sigma Aldrich, Cat No. 71380) was placed into the RSH tray. Each sample was measured in triplicates as well as additional Air, Buffer and Internal Standard (of each sample type spiked with 10 µL D-Hexanal stock (50 µL/L, >98 % purity, Merck, Cat No. 732338-250MG)) as controls (209 vials total). Each vial was incubated at 60 °C for 5 min, while the agitator was set to on for 5 seconds intervals. The headspace extraction of the sample was subsequently carried out with a Smart SPME Fiber (PAL System Smart SPME Fiber DVB/C-WR/PDMS (Divinylbenzene/Carbon Wide Range/Polydimethylsiloxane), phase 80 µm, fibre thickness 50/30 µm, fibre length 10 mm, colour code dark grey) at 40 °C for 15 min. The needle depth in the vial was set to 25 mm. The adsorbed sample was then injected in the SSL inlet set at 240 °C, with a desorption time of 5 min and an injection depth of 40 mm. To avoid carrying over the SPME-Arrow fibre was pre-conditioned in a dedicated conditioning station (PTV – Back Inlet) for 1 min at 240 °C. GC separation was carried out on a fused silica column (Agilent, DB-WAX GC Column, length 30 m, film thickness 0.25 mm, inner diameter 0.25 µm, format 7 inch cage, Cat No. 122-7032). Helium was the carrier gas used with a constant flow rate of 1.5 mL/min. The inlet temperature was set at 240 °C and with a split ratio of 6.67 (split flow 10 mL/min, spitless time 5 min). A ramped oven program was used, with an initial temperature of 40 °C holding for 3 min, followed by a ramp of 6 °C/min to reach 170 °C . A second ramp was set with 11 °C/min to reach a final temperature of 240 °C with a hold time of 3 min (to completely clean the column). The total run time was 35.1 min and the total analysis time was set to 43 min.

Generated GC-MS data was analysed in Chromeleon™ Chromatography Data System (CDS) Software (version 7.2). We used the Cobra peak detection algorithm (standard parameters) to integrate peaks and first compared all retrieved peaks (match factor 850) to the Wiley Registry of Mass Spectral Data (ISBN 978-1-119-73632-5) containing over 873,000 reference GC-MS spectra as well as the FFNSC3 (Flavors and Fragrances of Natural and Synthetic Compounds, ISBN: 978-1-119-06964-5) reference library of over 3000 mass spectra. Noise filtering was set to a relative threshold of 5 % and we removed all compounds before retention time (RT) 7-8 min as the eluting ethanol is skewing the spectra in this area as well as compounds with a RT > 32 min as these are mostly column contaminations. The confirming peaks (fragments) were manually adjusted according to the assigned compound to update the ion ratio and get more accurate total ion current (TIC) peaks. After an initial automated annotation based on retention times and fragmentation, MS reference spectra were added to each peak and peak annotation performed on a quality MS match within a retention window to provide more security against false negative annotations. Ultimately the generated TIC counts were exported, and the median per sample calculated and normalised using Total Sum Normalisation [37]. The Wiley [38], NIST

[39], and FFNSC [40] libraries were used to identify volatile compounds, yielding a total of 151 uniquely annotated volatiles.

**10. Untargeted Metabolomics with LC-MS**

Untargeted metabolomic analysis of microvinification samples was performed using high-resolution liquid chromatography-mass spectrometry (LC-MS) in both positive and negative ionization modes. LC-MS analyses were conducted using an Orbitrap mass analyzer operating in data-dependent acquisition (DDA) mode, with a mass range of 50–1500 m/z and mass resolutions of 60,000 at 200 m/z for MS1 and 30,000 at 200 m/z for MS2. Both positive and negative ionization modes were utilized, with stepped collision energies of 10, 30, and 45 NCE. Chromatographic separation was performed using a C18 column (e.g., Acquity BEH C18, 2.1 × 50 mm, 1.7 µm) at a flow rate of 300 µL/min and a temperature of 40 °C. The mobile phases consisted of 0.1 % formic acid in water (A) and 0.1 % formic acid in acetonitrile (B), with gradient and injection volume parameters specified based on the experimental setup.

Raw LC-MS data were processed with retention time alignment using ChromAlign, followed by unknown compound detection with a mass tolerance of 5 ppm, a minimum peak intensity threshold of 10,000, and a minimum of six scans per peak. Peak detection parameters included a chromatographic signal-to-noise threshold of 1.5, a peak width maximum of 0.1 minutes, and a gap ratio threshold of 0.35. Compound grouping was performed with a 5 ppm mass tolerance and a 0.05-minute retention time tolerance, using peak quality assessment that weighted area (3), coefficient of variation (10), and peak shape parameters including jaggedness, modality, and zig-zag index (5 each). Chemical background compounds were identified and hidden based on a comparison of sample-to-blank ratios. For normalization, areas were adjusted using constant median normalization, with blanks excluded from the calculations.

Compound annotation was performed with multiple databases, including mzCloud [41], mzVault [42], ChemSpider [43] with strict mass tolerance thresholds of 5–10 ppm for precursor and fragment ions. Spectral distance and match factor scoring were utilized to assess annotation confidence. Fragment ion annotations were refined through recalibrated libraries and ranked by similarity scores. Database searches for compound identification included BioCyc [44], the Human Metabolome Database [45], KEGG [46], Phenol-Explorer [47], PlantCyc [48], Arita Lab 6549 flavonoid structure database [49], and Natural Products Atlas masslist [50]. ChemSpider searches were performed using predicted compositions, with each compound limited to a maximum of three predicted compositions. Mass tolerance for ChemSpider searches was set to 5 ppm. mzCloud similarity searches utilized cosine matching algorithms to compare ddMS2 spectra, with similarity thresholds set to 50%. Mass range and resolution parameters were adjusted dynamically for accurate spectral interpretation, and isotopic patterns were integrated for precursor evaluation. In addition to compound detection, the analysis included gap-filling across all samples and compound ranking using the mzLogic algorithm [51]. This process utilized chemical background removal and normalization based on the constant median method, excluding blank samples from the calculations. The analysis was further supported by mapping compounds to biological pathways using the Metabolika module.

*Methodology Table 2. Total number of features identified by each mode of ionization, as well as number of MS1/M2 annotated features and MS2 annotated features.*

| **Mode** | **Features** | **Annotated Features: MS1/MS2** | **Annotated Features: MS2** |
|---|---|---|---|
| Negative | 5355 | 1389 | 170 |
| Positive | 6205 | 1942 | 191 |

For subsequent multi-omics analysis we used the MS2 identified features, and normalized tables with log transformation and Z-score normalization. The annotation accuracy of metabolites showing strong correlations (|corr| > 0.9) was further assessed to determine whether a metabolite could be an isomer, based on full matches to the predicted composition, mzCloud search, mzVault search, ChemSpider search, and the mass list search. Potential contamination was also evaluated by examining the sample-to-blank ratio (see Methodology Table 3).

*Methodology Table 3. LC-MS identified metabolites that either show strong correlations or rank among the top loadings in the variable plot. Metabolites were curated based on identification by at least two annotation sources, a high mzCloud match score (>70%), and a low likelihood of contamination (sample-to-blank ratio < 5).*

| **Name** | **Annot. Source: Predicted Compositions** | **Annot. Source: mzCloud Search** | **Annot. Source: mzVault Search** | **Annot. Source: ChemSpider Search** | **Annot. Source: MassList Search** | **mzCloud Best Match** | **mzCloud Best Match Confidence** | **Ratio: Sample / Blank** |
|---|---|---|---|---|---|---|---|---|
| **5-Hydroxytryptophol** | Full match | No results | Full match | Partial match | Partial match | NaN | NaN | 1.8 |
| **Azelaic acid** | Full match | Full match | Full match | Full match | Full match | 99.1 | 10.0 | 0.8 |
| **Norepinephrine** | Full match | Full match | Full match | Not the top hit | Not the top hit | 88.7 | 55.2 | 4.6 |
| **N-Methyldioctylamine** | Full match | Full match | No results | No results | No results | 79.6 | 49.8 | 0.4 |
| **4-Acetamidobutanoic acid** | Full match | Full match | No results | Full match | Not the top hit | 78.8 | 8.9 | 2.8 |
| **N-(5-acetamidopentyl)acetamide** | Full match | Full match | No results | Partial match | No results | 72.0 | 50.8 | 0.4 |
| **Hexadecanedioic acid** | Full match | Full match | No results | Full match | Full match | 71.2 | 8.6 | 2.0 |
| **C16-0(Palmitoyl)ceramide** | Full match | No results | Full match | No results | Partial match | NaN | NaN | 1.7 |

| | | | | | | | | |
|---|---|---|---|---|---|---|---|---|
| **Phenylalanyltyrosine** | Full match | No results | Full match | Not the top hit | Partial match | NaN | NaN | 2.3 |
| **Hydroxyprolyl-Proline** | Full match | No results | Full match | Partial match | Partial match | NaN | NaN | 4.8 |
| **Pro-Phe** | Not the top hit | No results | Full match | Partial match | Partial match | NaN | NaN | 1.7 |
| **1-Palmitoyl-sn-glycero-3-phosphocholine** | Not the top hit | No results | Full match | Partial match | Full match | NaN | NaN | 0.3 |
| **L-Glutamic acid** | Full match | Full match | Full match | Full match | Not the top hit | 86.4 | 70.6 | 0.3 |
| **Decaethyleneglycol** | Not the top hit | Full match | No results | No results | No results | 77.3 | 73.2 | 1.0 |
| **Undecaethylene glycol** | Not the top hit | Full match | No results | No results | No match | 75.4 | 71.8 | 1.0 |
| **Palmitoyl ethanolamide** | Full match | Full match | No results | Full match | Full match | 88.3 | 9.4 | 0.3 |
| **2-Amino-1,3,4-octadecanetriol** | Full match | Full match | Full match | Full match | Full match | 79.8 | 9.0 | 1.4 |
| **γ-L-glutaminyl-3,4-benzoquinone** | Full match | Full match | No results | Partial match | Not the top hit | 84.8 | 9.2 | 0.9 |

14. Callahan BJ, Sankaran K, Fukuyama JA *et al.* Bioconductor Workflow for Microbiome Data

Analysis: from raw reads to community analyses. *F1000Research* 2016;**5**:1492.
15. Bokulich NA, Kaehler BD, Rideout JR *et al.* Optimizing taxonomic classification of marker-gene amplicon sequences with QIIME 2's q2-feature-classifier plugin. *Microbiome* 2018;**6**:90.
16. Quast C, Pruesse E, Yilmaz P *et al.* The SILVA ribosomal RNA gene database project: improved data processing and web-based tools. *Nucleic Acids Res* 2012;**41**:D590–6.
17. Ii MSR, O'Rourke DR, Kaehler BD *et al.* RESCRIPt: Reproducible sequence taxonomy reference database management. *PLOS Comput Biol* 2021;**17**:e1009581.
18. Kaehler BD, Bokulich NA, McDonald D *et al.* Species abundance information improves sequence taxonomy classification accuracy. *Nat Commun* 2019;**10**:4643.
19. Abarenkov K, Zirk A, Piirmann T *et al.* UNITE QIIME release for Fungi. 2023, DOI: 10.15156/BIO/2938079.
20. Pielou EC. The measurement of diversity in different types of biological collections. *J Theor Biol* 1966;**13**:131–44.
21. Shannon CE. A Mathematical Theory of Communication. *Bell Syst Tech J* 1948;**27**:379–423.
22. Sorensen Thorvald. A method of establishing groups of equal amplitude in plant sociology based on similarity of species content and its application to analyses of the vegetation on Danish commons. *Biol Skr* 1948;**5**:1–34.
23. Jaccard P. Nouvelles recherches sur la distribution florale. *Bull Soc Vaud Sci Nat* 1908;**44**:223–70.
24. Bokulich NA. Integrating sequence composition information into microbial diversity analyses with k-mer frequency counting. Rosen G (ed.). *mSystems* 2025:e01550-24.
25. Price MN, Dehal PS, Arkin AP. FastTree 2 – Approximately Maximum-Likelihood Trees for Large Alignments. Poon AFY (ed.). *PLoS ONE* 2010;**5**:e9490.
26. Katoh K, Standley DM. MAFFT Multiple Sequence Alignment Software Version 7: Improvements in Performance and Usability. *Mol Biol Evol* 2013;**30**:772–80.
27. Lozupone C, Lladser ME, Knights D *et al.* UniFrac: an effective distance metric for microbial community comparison. *ISME J* 2011;**5**:169–72.
28. Anderson MJ. A new method for non-parametric multivariate analysis of variance. *Austral Ecol* 2001;**26**:32–46.
29. Oksanen J, Simpson GL, Blanchet FG *et al.* vegan: Community Ecology Package. 2001:2.6-8.
30. Halford M. Prince. 2025, DOI: https://github.com/MaxHalford/prince.
31. Rohart F, Gautier B, Singh A *et al.* mixOmics: An R package for 'omics feature selection and

multiple data integration. Schneidman D (ed.). *PLOS Comput Biol* 2017;**13**:e1005752.
32. Foster ZSL, Sharpton TJ, Grünwald NJ. Metacoder: An R package for visualization and manipulation of community taxonomic diversity data. Poisot T (ed.). *PLOS Comput Biol* 2017;**13**:e1005404.
33. Lin H, Peddada SD. Multigroup analysis of compositions of microbiomes with covariate adjustments and repeated measures. *Nat Methods* 2024;**21**:83–91.
34. The Matplotlib Development Team. Matplotlib: Visualization with Python. 2024, DOI: 10.5281/ZENODO.13308876.
35. Waskom M. seaborn: statistical data visualization. *J Open Source Softw* 2021;**6**:3021.
36. Jordahl K, Bossche JVD, Fleischmann M *et al.* geopandas/geopandas: v0.8.1. 2020, DOI: 10.5281/ZENODO.3946761.
37. Noonan MJ, Tinnesand HV, Buesching CD. Normalizing Gas-Chromatography–Mass Spectrometry Data: Method Choice can Alter Biological Inference. *BioEssays* 2018;**40**:1700210.
38. Wiley. Wiley Registry of Mass Spectral Data, 10th Edition.
39. National Institute of Standards and Technology. NIST Mass Spectral Library (Version).
40. Mondello, Luigi. Wiley FFNSC Library - Mass Spectra of Flavors and Fragrances of Natural and Synthetic Compounds, 3rd Edition. 2015.
41. HighChem LLC. mzCloud - Advanced Mass Spectral Database.
42. Thermo Fisher Scientific Inc. mzVault.
43. Royal Society of Chemistry. ChemSpider. 2025.
44. Bioinformatics Research Group, SRI International. BioCyc.
45. Wishart DS, Tzur D, Knox C *et al.* HMDB: the Human Metabolome Database. *Nucleic Acids Res* 2007;**35**:D521–6.
46. Kanehisa M, Furumichi M, Sato Y *et al.* KEGG: biological systems database as a model of the real world. *Nucleic Acids Res* 2025;**53**:D672–7.
47. Neveu V, Perez-Jimenez J, Vos F *et al.* Phenol-Explorer: an online comprehensive database on polyphenol contents in foods. *Database* 2010;**2010**:bap024–bap024.
48. Hawkins C, Xue B, Yasmin F *et al.* Plant Metabolic Network 16: expansion of underrepresented plant groups and experimentally supported enzyme data. *Nucleic Acids Res* 2025;**53**:D1606–13.
49. Arita M, Suwa K. Search extension transforms Wiki into a relational system: A case for flavonoid metabolite database. *BioData Min* 2008;**1**:7.
50. Poynton EF, van Santen JA, Pin M *et al.* The Natural Products Atlas 3.0: extending the database of microbially derived natural products. *Nucleic Acids Res* 2025;**53**:D691–9.

51. Thermo Fisher Scientific Inc. mzLogic Data Analysis Algorithm: Accelerate small-molecule unknown identification. 2019.
