## Supplementary Figures and Tables for "Grape Expectations: Disentangling Environmental Drivers of Microbiome Establishment in Winegrowing Ecosystems"

### Supplementary Tables

***Supplementary Table 1.*** *Spatiotemporal variation in alpha diversity (evenness, observed features, Shannon entropy) of bacterial communities across different sample types.*

| | Soil | | | Bark | | |
|---|---|---|---|---|---|---|
| | **Evenness** | **Observed Features** | **Shannon** | **Evenness** | **Observed Features** | **Shannon** |
| Time Point | 0.6334 | 0.3031 | 0.2981 | | | |
| Plot | 0.0793 | 0.2772 | 0.1119 | 0.1198 | 0.2166 | 0.0917 |
| Year | 0.0050 | 0.0037 | 0.0022 | 0.8889 | 0.0029 | 0.2143 |

***Supplementary Table 2****. Spatiotemporal variation in alpha diversity (evenness, observed features, Shannon entropy) of fungal communities across different sample types.*

| | Soil | | | Berries | | | Bark | | | Leaves | | |
|---|---|---|---|---|---|---|---|---|---|---|---|---|
| | **Evenness** | **Observed Features** | **Shannon** | **Evenness** | **Observed Features** | **Shannon** | **Evenness** | **Observed Features** | **Shannon** | **Evenness** | **Observed Features** | **Shannon** |
| Time Point | 0.4450 | 0.1852 | 0.5995 | 0.1080 | 0.0002 | 0.6157 | | | | | | |
| Plot | 0.3487 | 0.4093 | 0.2358 | 0.0000 | 0.1082 | 0.0000 | 0.3822 | 0.5101 | 0.5753 | 0.8611 | 0.9999 | 0.8980 |
| Year | 0.9640 | 0.6017 | 0.5812 | 0.0637 | 0.0000 | 0.2143 | 0.8472 | 0.2568 | 0.3341 | 0.3858 | 0.0015 | 0.0280 |

***Supplementary Table 3****. Statistically testing distance decay relationships in soil using a Mantel test to assess correlations between beta diversity metrics and the geodesic distance between vineyards, calculated for all samples, as well as subset to individual time points or years.*

| | ITS | | | | | | 16S | | | |
|---|---|---|---|---|---|---|---|---|---|---|
| | **All soil samples** | | **2021 samples** | | **Veraison 2021** | | **All soil samples** | | **Spring soil** | |
| | **Mantel R** | **p-value** | **Mantel R** | **p-value** | **Mantel R** | **p-value** | **Mantel R** | **p-value** | **Mantel R** | **p-value** |
| Bray Curtis | 0.040 | 0.300 | -0.122 | 0.656 | -0.100 | 0.606 | -0.055 | 0.778 | -0.165 | 0.881 |
| Jaccard | 0.059 | 0.202 | -0.187 | 0.737 | -0.207 | 0.775 | -0.054 | 0.750 | -0.083 | 0.686 |
| weighted UniFrac | | | | | | | -0.074 | 0.859 | -0.070 | 0.668 |
| unweighted UniFrac | | | | | | | -0.100 | 0.942 | -0.274 | 0.988 |

| | | | | | | | | | | |
|---|---|---|---|---|---|---|---|---|---|---|
| Bray Curtis (kmer) | 0.098 | 0.116 | 0.192 | 0.201 | 0.183 | 0.238 | -0.086 | 0.901 | -0.264 | 0.981 |
| Jaccard (kmer) | 0.134 | 0.050 | -0.037 | 0.483 | 0.046 | 0.381 | -0.042 | 0.716 | -0.058 | 0.673 |

***Supplementary Table 4****. Statistically testing distance decay relationships in bark using a Mantel test to assess correlations between beta diversity metrics and the geodesic distance between vineyards using a  Mantel Test of soil samples, calculated for all samples across all time points, as well as subset to individual time points or years.*

| | **ITS** | | | | **16S** | |
|---|---|---|---|---|---|---|
| | **All soil samples** | | **Veraison 2021 samples** | | **All soil samples** | |
| | **Mantel R** | **p-value** | **Mantel R** | **p-value** | **Mantel R** | **p-value** |
| Bray Curtis | 0.071 | 0.179 | 0.007 | 0.482 | -0.003 | 0.470 |
| Jaccard | 0.021 | 0.386 | -0.002 | 0.493 | 0.010 | 0.420 |
| weighted UniFrac | | | | | -0.016 | 0.517 |
| unweighted UniFrac | | | | | 0.075 | 0.258 |
| Bray Curtis (kmer) | 0.090 | 0.116 | 0.073 | 0.355 | 0.024 | 0.361 |
| Jaccard (kmer) | 0.069 | 0.201 | -0.003 | 0.440 | 0.012 | 0.452 |

**Supplementary Table 5.** Predicting the vineyard from one year for the other using Random Forest machine learning.

| | | | | |
|---|---|---|---|---|
| Training Year | 2021 | 2021 | 2023 | 2022 |
| Training Sample Number | 193 | 193 | 50 | 50 |
| Training % | 79.42 | 79.42 | 50 | 50 |
| Test Year | 2022 | 2023 | 2022 | 2023 |
| Test Sample Number | 50 | 50 | 50 | 50 |
| Test % | 20.58 | 20.58 | 50 | 50 |
| **ROC Mean AUC** | **0.74** | **0.62** | **0.76** | **0.74** |

***Supplementary Table 6***. *Statistically testing distance decay relationships in berry samples of Appellation d'origine contrôlée (AOC) Lavaux using a Mantel test to assess correlations between beta diversity metrics and the geodesic distance between vineyards, calculated for all samples across all time points, as well as subset of the densely sampled 2021 harvest.*

| | **All harvest samples** | | **Harvest 2021** | |
|---|---|---|---|---|
| | **Mantel R** | **p-value** | **Mantel R** | **p-value** |
| Bray Curtis | 0.195 | 0.001 | 0.279 | 0.001 |
| Jaccard | 0.176 | 0.001 | 0.212 | 0.001 |
| Bray Curtis (kmer) | 0.189 | 0.001 | 0.270 | 0.001 |
| Jaccard (kmer) | 0.054 | 0.019 | 0.102 | 0.002 |

***Supplementary Table 7***. *Statistically testing distance decay relationships in berry samples of different vines within the same vineyards using a Mantel test.*

| | **Plot_4** | | **Plot_5** | | **Plot_9** | | **Plot_11** | | **Plot_12** | | **Plot_13** | |
|---|---|---|---|---|---|---|---|---|---|---|---|---|
| | **Mantel R** | **p-value** | **Mantel R** | **p-value** | **Mantel R** | **p-value** | **Mantel R** | **p-value** | **Mantel R** | **p-value** | **Mantel R** | **p-value** |
| Bray Curtis | -0.015 | 0.531 | 0.033 | 0.378 | 0.117 | 0.101 | -0.105 | 0.900 | -0.045 | 0.647 | 0.113 | 0.263 |
| Jaccard | -0.039 | 0.712 | -0.169 | 0.916 | 0.030 | 0.378 | -0.191 | 0.994 | -0.056 | 0.672 | 0.272 | 0.055 |
| Bray Curtis (kmer) | -0.019 | 0.559 | 0.075 | 0.262 | 0.031 | 0.338 | -0.057 | 0.736 | -0.009 | 0.493 | 0.236 | 0.075 |
| Jaccard (kmer) | -0.081 | 0.837 | -0.241 | 0.999 | -0.083 | 0.836 | -0.018 | 0.571 | -0.076 | 0.705 | 0.073 | 0.334 |

| | **Plot_14** | | **Plot_15** | | **Plot_18** | | **Plot_19** | | **Plot_20** | | **Plot_17** | |
|---|---|---|---|---|---|---|---|---|---|---|---|---|
| | **Mantel R** | **p-value** | **Mantel R** | **p-value** | **Mantel R** | **p-value** | **Mantel R** | **p-value** | **Mantel R** | **p-value** | **Mantel R** | **p-value** |
| Bray Curtis | 0.345 | 0.001 | -0.060 | 0.769 | -0.198 | 0.836 | 0.486 | 0.167 | 0.036 | 0.383 | 0.065 | 0.105 |
| Jaccard | 0.222 | 0.027 | 0.052 | 0.216 | -0.069 | 0.623 | -0.714 | 1.000 | 0.009 | 0.469 | 0.058 | 0.142 |
| Bray Curtis (kmer) | 0.308 | 0.010 | -0.012 | 0.536 | -0.165 | 0.785 | 0.486 | 0.167 | -0.086 | 0.789 | 0.091 | 0.060 |
| Jaccard (kmer) | 0.053 | 0.285 | 0.102 | 0.075 | -0.131 | 0.756 | 0.143 | 0.500 | 0.137 | 0.101 | 0.116 | 0.031 |

***Supplementary Table 8***. *Statistically testing distance decay relationships in berry samples of Appellation d'origine contrôlée (AOC) Lavaux as well as AOC Valais using a Mantel test to assess correlations between beta diversity metrics and the geodesic distance between vineyards, calculated for all samples across harvest samples as well as subset for the separate varieties.*

| **Lavaux & Valais 2023 harvest samples** | **Lavaux & Valais 2023 harvest samples** | **Lavaux & Valais 2023 harvest samples** |
|---|---|---|

| | | | (only Chasselas) | | (only Pinot Noir) | |
|---|---|---|---|---|---|---|
| | **Mantel R** | **p-value** | **Mantel R** | **p-value** | **Mantel R** | **p-value** |
| Bray Curtis | 0.422 | 0.001 | 0.624 | 0.001 | 0.125 | 0.017 |
| Jaccard | 0.421 | 0.001 | 0.317 | 0.001 | 0.156 | 0.006 |
| Bray Curtis (kmer) | 0.465 | 0.001 | 0.599 | 0.001 | 0.162 | 0.005 |
| Jaccard (kmer) | 0.445 | 0.001 | 0.284 | 0.001 | 0.192 | 0.003 |

***Supplementary Table 9**. Permutational analysis of variance (PERMANOVA) showing the relationship between beta diversity of berry fungal communities and climate variables, including median relative humidity (RH) and median temperature.*

| | **Bray Curtis** | | **Jaccard** | | **Kmer: Bray Curtis** | | **Kmer: Jaccard** | |
|---|---|---|---|---|---|---|---|---|
| | **R2** | **p-val** | **R2** | **p-val** | **R2** | **p-val** | **R2** | **p-val** |
| **Median RH** | 0.022 | 0.001 | 0.015 | 0.001 | 0.042 | 0.001 | 0.027 | 0.001 |
| **Median Temperature** | 0.004 | 0.164 | 0.005 | 0.001 | 0.006 | 0.098 | 0.006 | 0.003 |
| **Residuals** | 0.974 | NaN | 0.980 | NaN | 0.953 | NaN | 0.967 | NaN |

***Supplementary Table 10**. Permutational analysis of variance (PERMANOVA) showing the relationship between beta diversity of berry fungal communities and climate variables when nested for year.*

| | **Bray Curtis** | | **Jaccard** | | **Kmer: Bray Curtis** | | **Kmer: Jaccard** | |
|---|---|---|---|---|---|---|---|---|
| | **R2** | **p-val** | **R2** | **p-val** | **R2** | **p-val** | **R2** | **p-val** |
| **Year : median RH** | 0.022 | 0.001 | 0.015 | 0.001 | 0.042 | 0.001 | 0.027 | 0.001 |
| **Year : median temperature** | 0.004 | 0.157 | 0.005 | 0.001 | 0.006 | 0.101 | 0.006 | 0.002 |
| **Residuals** | 0.974 | NaN | 0.980 | NaN | 0.953 | NaN | 0.967 | NaN |

***Supplementary Table 11**. Mantel test assessing the correlation between berry fungal community dissimilarities and Euclidean distances of median temperature and median relative humidity.*

| | Climate: median temp & median rh | | Temperature: median, min, max | | Relative humidity: median, min, max | |
|---|---|---|---|---|---|---|
| | Spearman rho | p-value | Spearman rho | p-value | Spearman rho | p-value |
| Bray Curtis | -0.058 | 0.972 | -0.028 | 0.809 | -0.060 | 0.981 |
| Jaccard | -0.089 | 1.000 | -0.096 | 1.000 | -0.111 | 1.000 |
| Bray Curtis (kmer) | 0.016 | 0.257 | 0.052 | 0.044 | 0.017 | 0.288 |
| Jaccard (kmer) | 0.332 | 0.001 | 0.345 | 0.001 | 0.323 | 0.001 |

***Supplementary Table 12.*** *Strongest correlations between metabolites and microbes (|corr| > 0.9) identified by the DIABLO model (Data Integration Analysis for Biomarker Discovery using a Latent Variable Approach for Omics studies). The table includes only LC-MS metabolites with high-confidence annotations and confirmed to not be potential contaminants (see Supplementary Methodology).*

| Feature 1 | Feature 2 | Correlation |
|---|---|---|
| Acetobacter pasteurianus | N-Methyldioctylamine | -0.936 |
| Acetobacter pasteurianus | trans-Aconitic acid | -0.922 |
| Acetobacter pasteurianus | Hexadecanedioic acid | -0.957 |
| Acetobacter pasteurianus | Saccharomyces cerevisiae | -0.994 |
| Benzaldehyde, 2,4-dimethyl- | Hydroxyprolyl-Proline | -0.914 |
| Diethyl succinate | Hydroxyprolyl-Proline | -0.911 |
| Diethyl succinate | Benzaldehyde, 2,4-dimethyl- | 0.955 |
| Diethyl succinate | Hexanal | 0.920 |
| Farnesol, (2Z,6Z)- | 5-Hydroxytryptophol | -0.935 |
| Hanseniaspora sp. | Farnesol, (2Z,6Z)- | 0.909 |
| Hanseniaspora sp. | Tetrahydro lavandulyl acetate | 0.945 |
| Hexadecanedioic acid | N-Methyldioctylamine | 0.940 |
| Hexadecanedioic acid | trans-Aconitic acid | 0.909 |
| Hexanal | Benzaldehyde, 2,4-dimethyl- | 0.922 |
| Mycosphaerella tassiana | 4-Acetamidobutanoic acid | -0.939 |

| | | |
|---|---|---|
| Mycosphaerella tassiana | 3-Hexen-1-ol, (Z)- | 0.930 |
| Mycosphaerella tassiana | Benzaldehyde, 2,4-dimethyl- | 0.930 |
| Mycosphaerella tassiana | Diethyl succinate | 0.923 |
| Mycosphaerella tassiana | Hexadecanoic acid, ethyl ester | -0.929 |
| Saccharomyces cerevisiae | Azelaic acid | 0.918 |
| Saccharomyces cerevisiae | Hexadecanedioic acid | 0.937 |
| Tetradecane | Benzaldehyde, 2,4-dimethyl- | 0.906 |
| Tetradecane | Diethyl succinate | 0.903 |
| Tetrahydro lavandulyl acetate | N-Methyldioctylamine | -0.972 |
| trans-Aconitic acid | N-Methyldioctylamine | 0.936 |

### Supplementary Figures

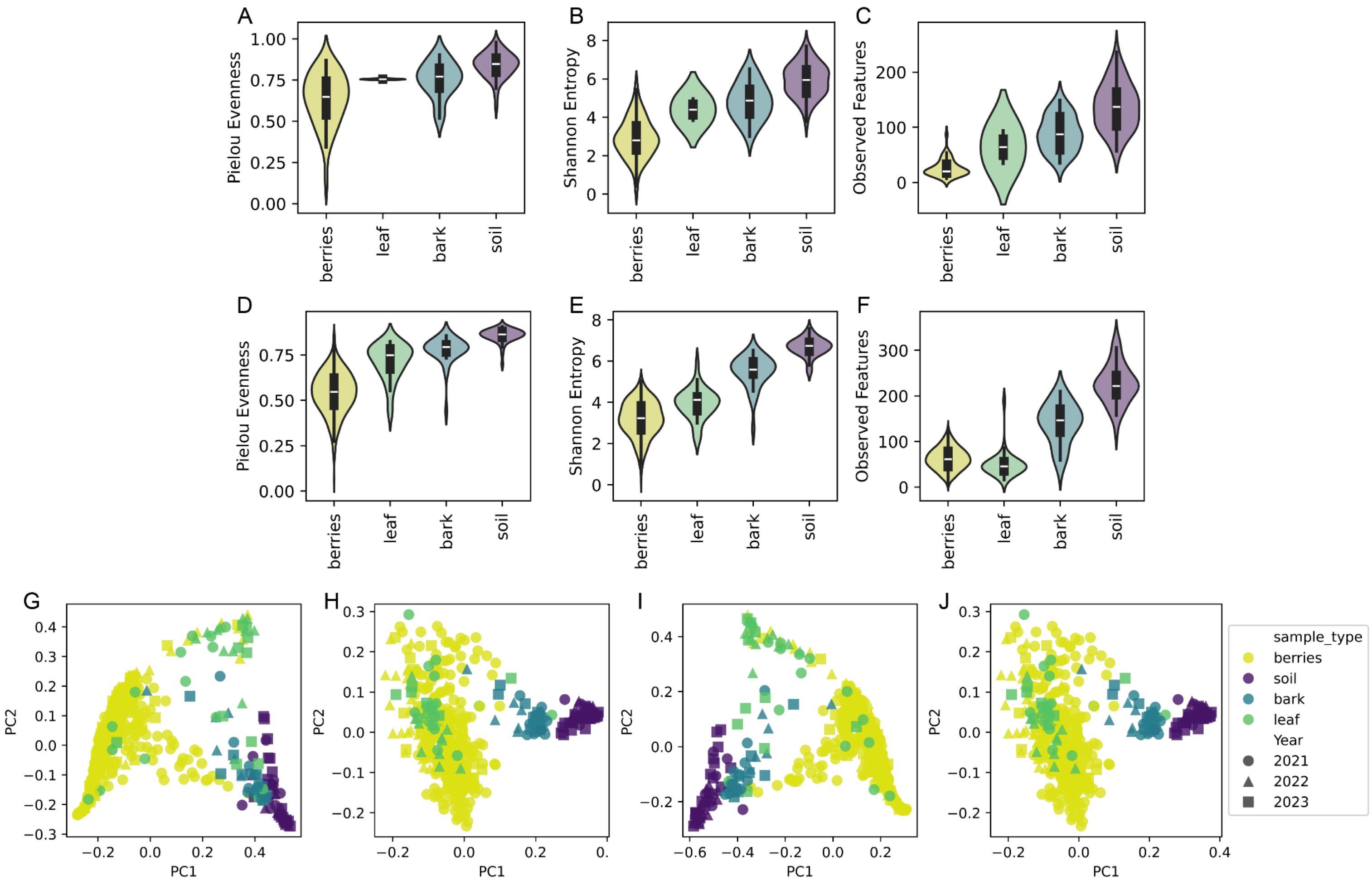


***Supplementary Figure 1.*** *Alpha and Beta diversity comparisons of microbial communities across soil, bark, leaves and berries (evenly rarefied across all sample types), for bacteria (A-C) and fungi (D-J). Showing various alpha diversity metrics: (A,D) Evenness, (B,E) Shannon Entropy, (C,F) Observed Features. Beta diversity of fungal communities are shown in PCoA plots, with points colored by sample type and shaped by year. Distance metrics include: (G) Bray-Curtis, (H) Jaccard, (I) k-mer-based Jaccard, and (J) k-mer-based Bray-Curtis.*

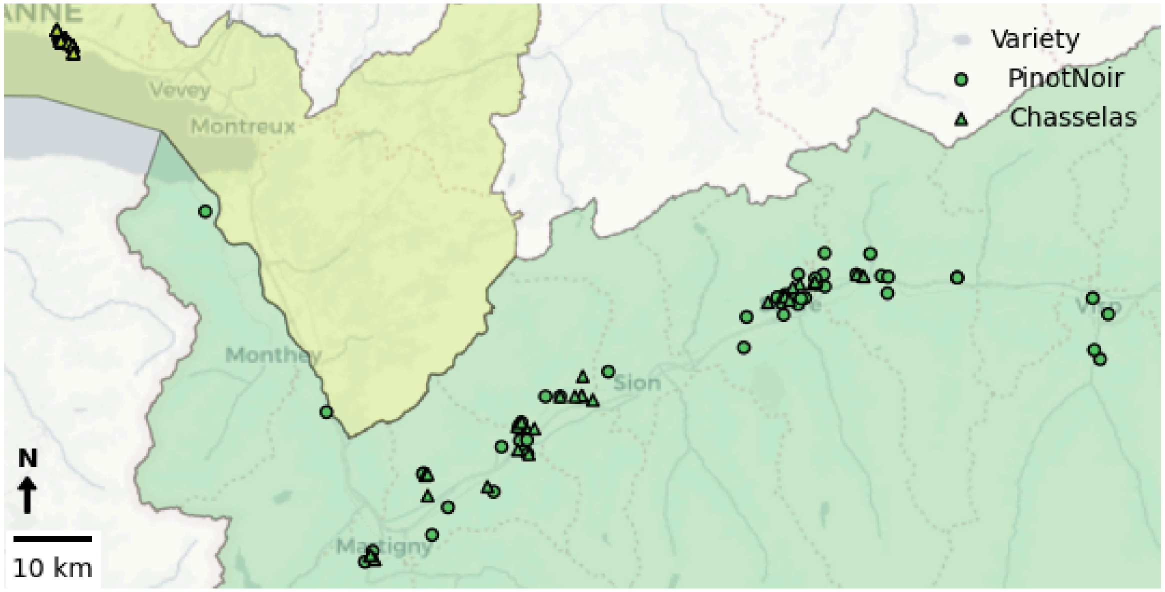


***Supplementary Figure 2.*** Map showing the samples collected from *Appellation d'origine contrôlée* (AOC) Lavaux in canton Vaud (light green) and from AOC Valais (dark green) from Pinot Noir (circle) and Chasselas (triangle) at harvest in 2023. The maximal distance between two vineyards is 95.21 km.

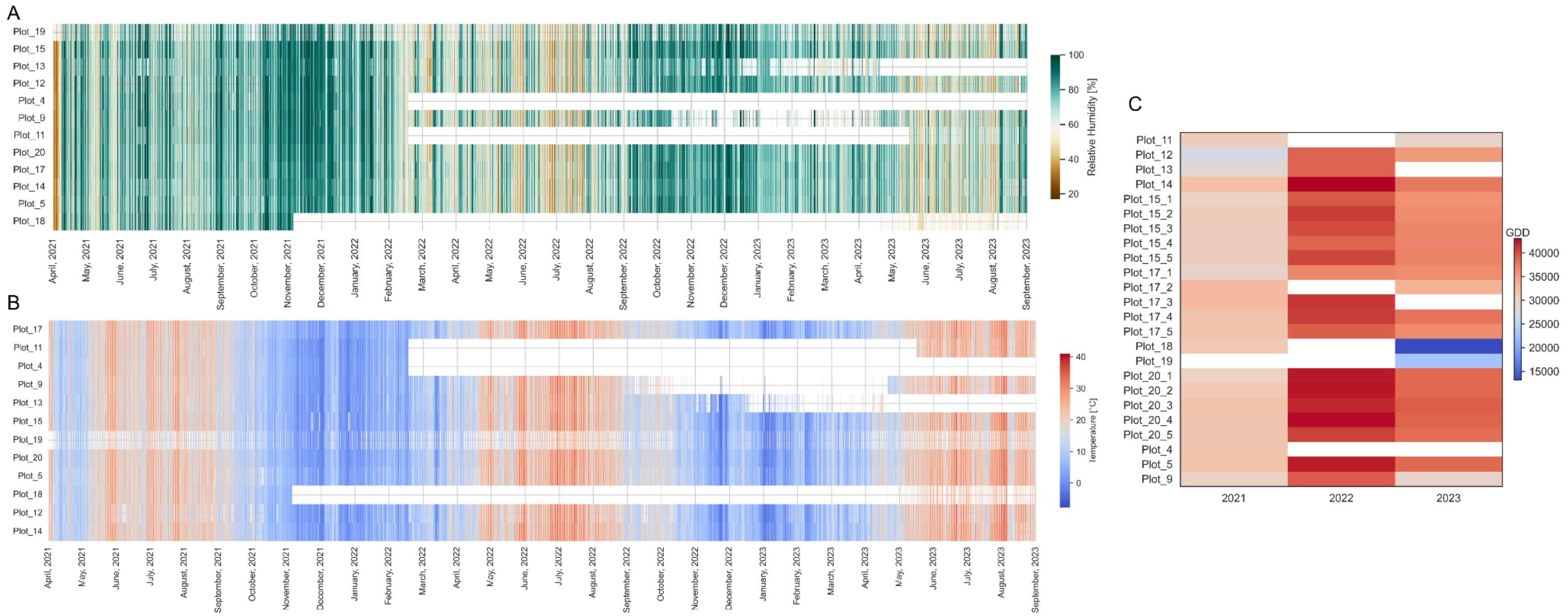


***Supplementary Figure 3.*** *Climatic variations of the Appellation d'origine contrôlée* (AOC) *Lavaux vineyards, as continuously measured by the installed sensors, showing the relative humidity (A) and temperature (B) of the plots, as well as calculated growing degree days (GDD) varying for each plot (D).*

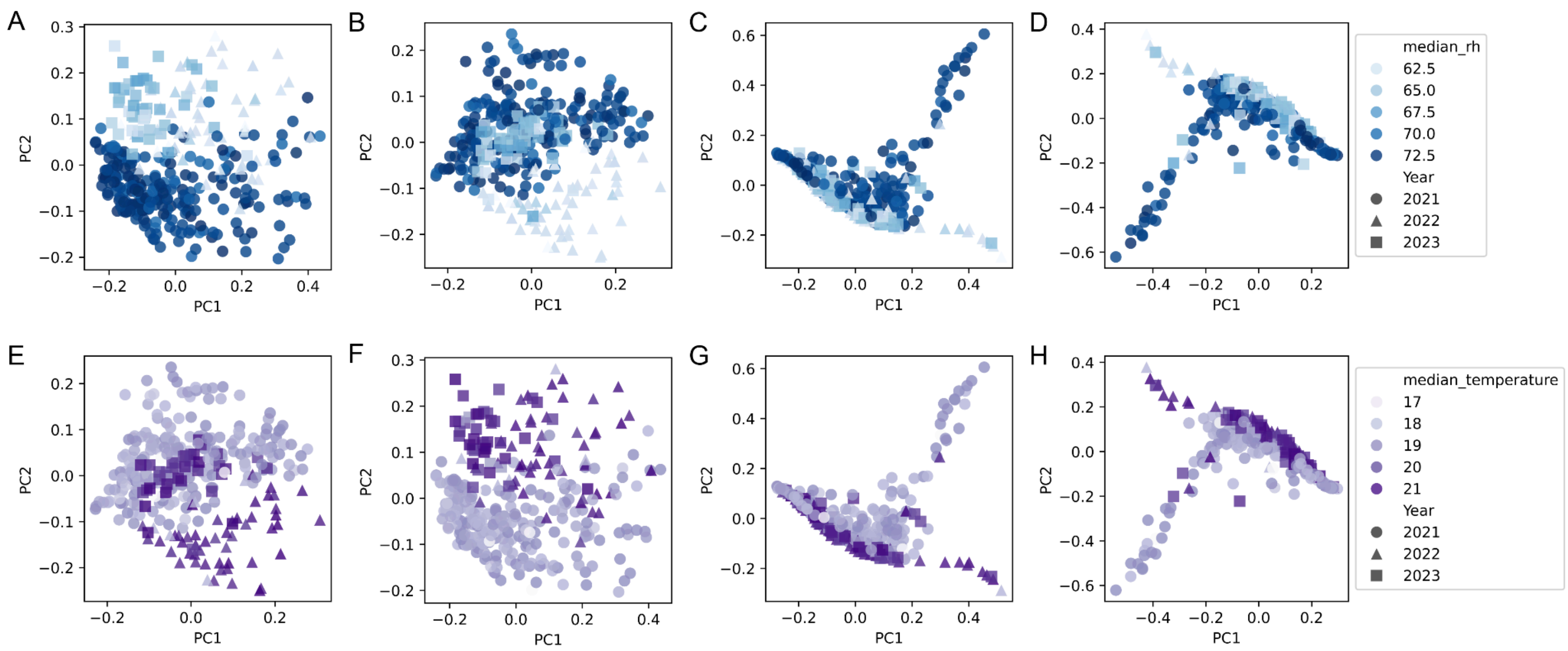


***Supplementary Figure 4.*** - PCoA plots of berry fungal communities, colored by relative humidity (A–D) and temperature (E–H), for various beta diversity metrics: (A, E) Jaccard, (B, F) k-mer-based Jaccard, (C, G) Bray-Curtis, and (D, H) k-mer-based Bray-Curtis.

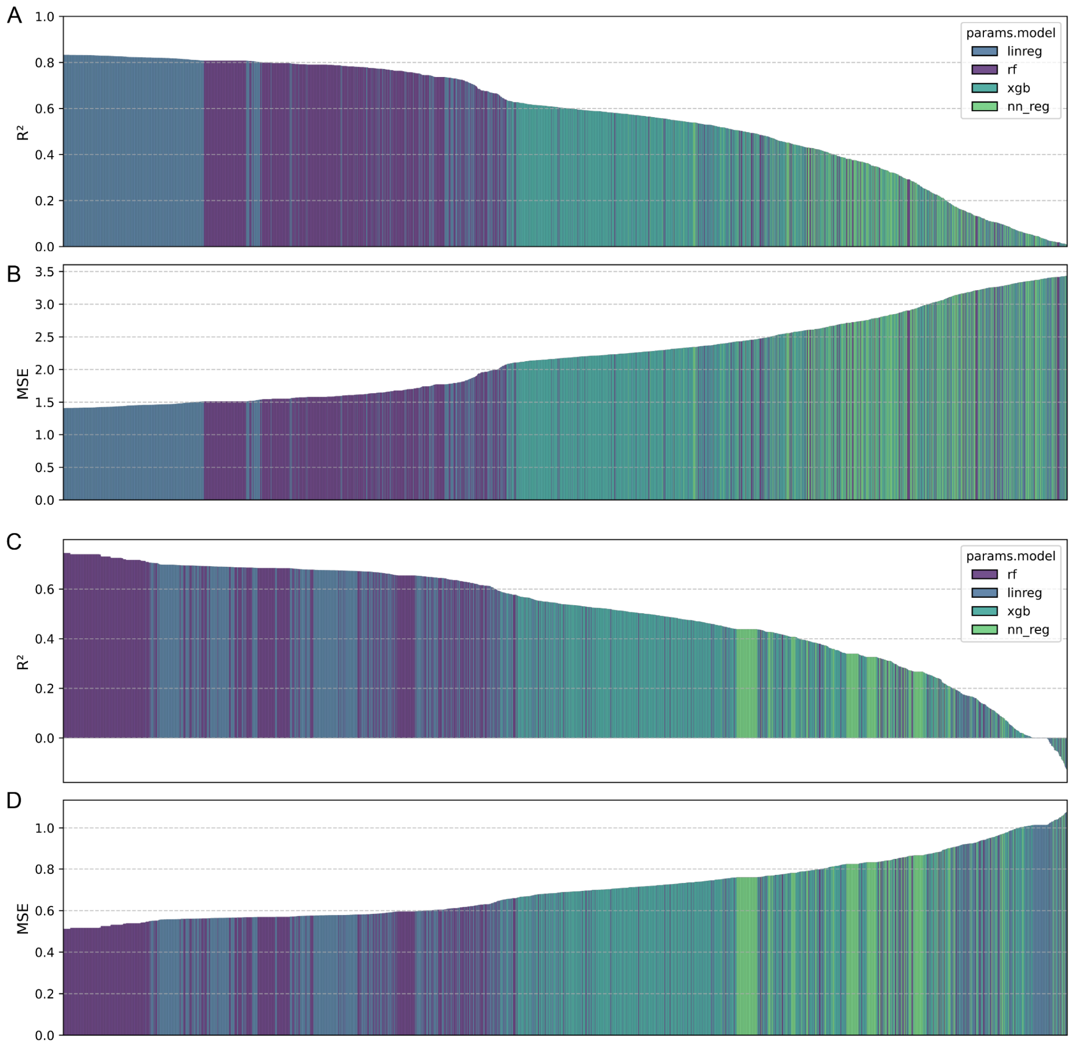


***Supplementary Figure 5.*** *Validation of the test set for 900 trials of the four models tested in RITME: linear regression (linreg), extreme gradient boosting (xgb), neural network regression (nn_reg), and random forest regression (rf). The framework applies various data transformations, agglomerations, feature selection methods, and optimizes for low mean square error (MSE, panels B and D) and high explained variance ($R^2$, panels A and C) in the test dataset. While all models are evaluated, the best-performing model is selected: linear regression for predicting relative humidity (A and B) and random forest for predicting temperature (C and D).*

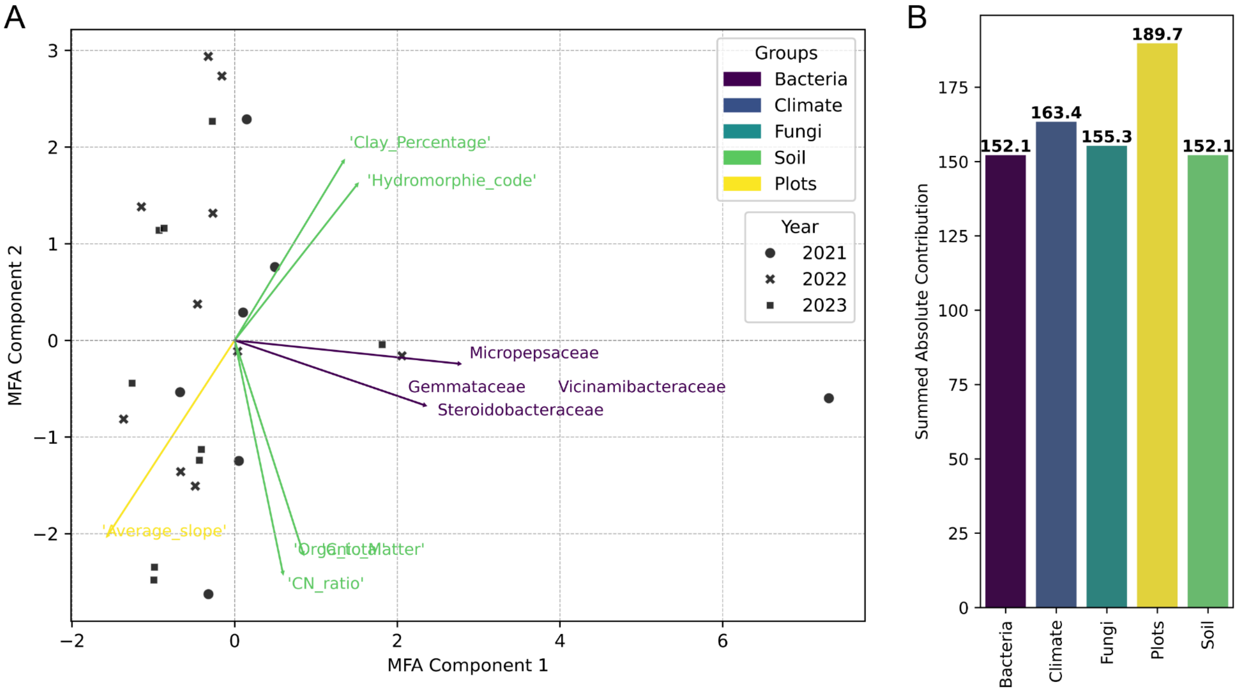


***Supplementary Figure 6.*** *Multifactor Analysis (MFA) of soil microbial communities and environmental factors, showing (A) the biplot with top 10 loadings and (B) the absolute group contributions.*

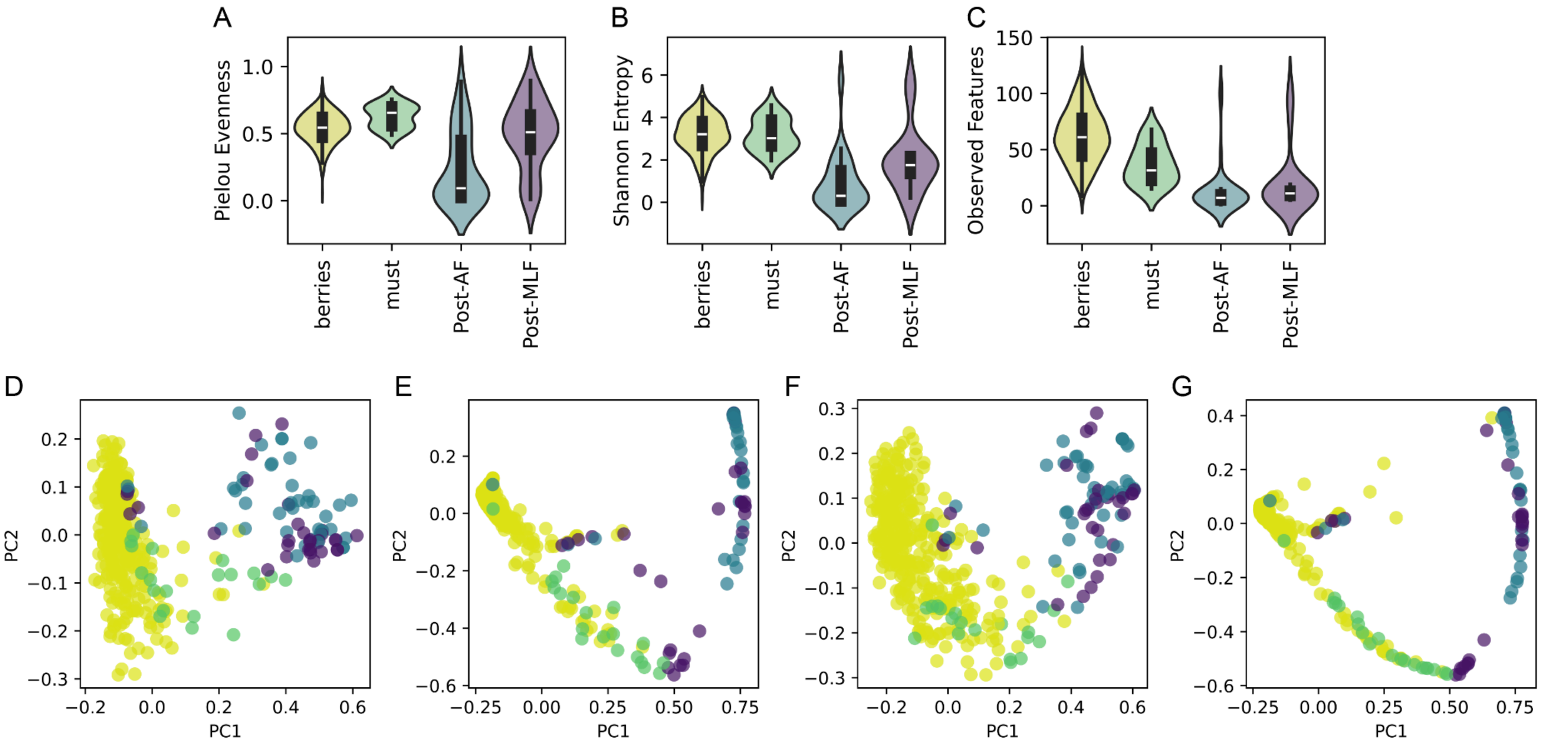


***Supplementary Figure 7***: *Alpha and beta diversity metrics illustrating microbial dynamics from fruit to wine. Alpha diversity is shown across different sample types: berries (yellow), pressed must (green), microvinification samples after initial alcoholic fermentation (Post-AF, blue), and final wine after malolactic fermentation (Post-MLF, purple), for: (A) Evenness, (B) Shannon Entropy, (C) Observed Features. Beta diversity is displayed in PCoA plots for these sample types using: (D) Jaccard, (E) Bray-Curtis, (F) k-mer based Jaccard, and (G) k-mer based Bray-Curtis.*

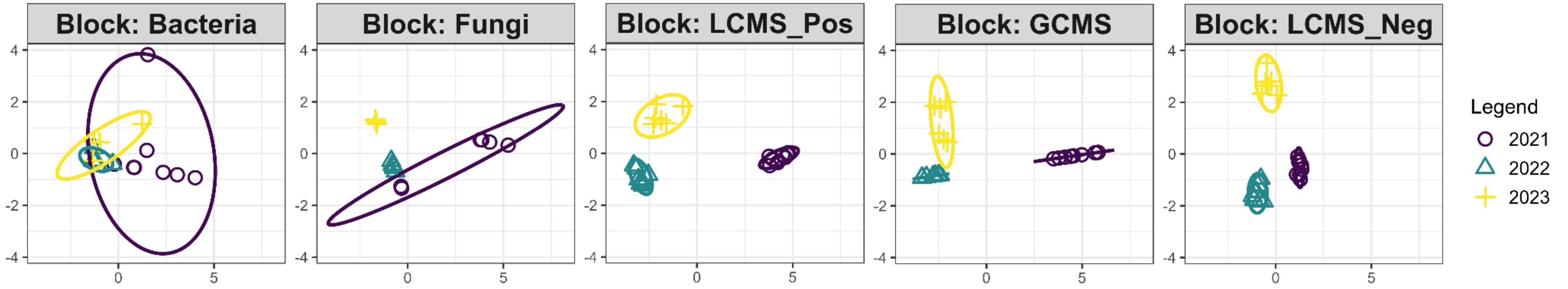


***Supplementary Figure 8.*** *Partial Least Squares (PLS) scores (variable 1 vs variable 2) of the DIABLO (Data Integration Analysis for Biomarker Discovery using a Latent Variable Approach for Omics studies) model integrating bacterial and fungal microbiome data with untargeted metabolomics data (GC-MS and LC-MS) show robust clustering per year for all datasets.*

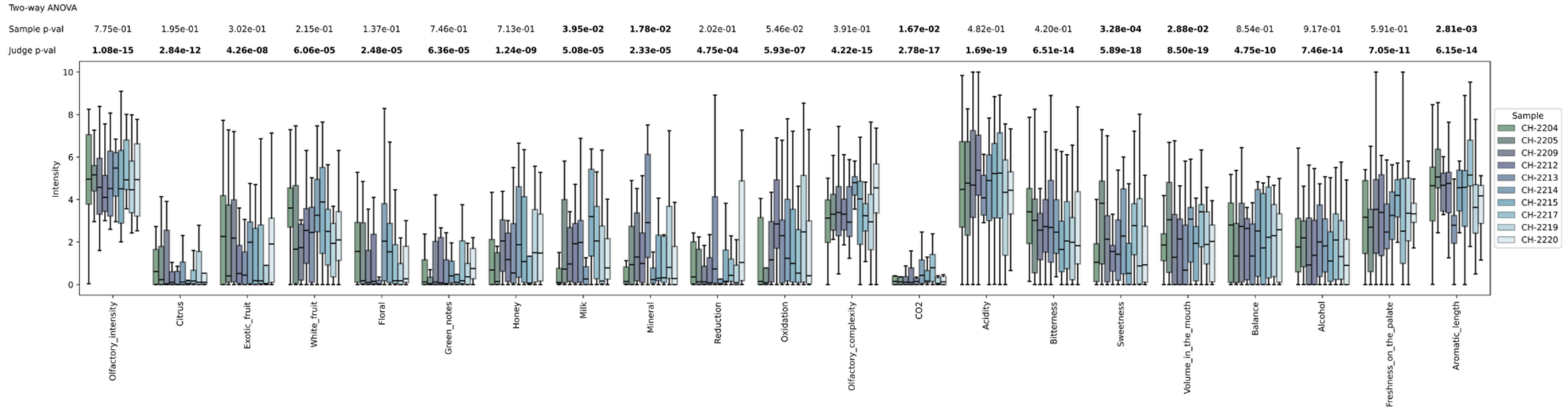


***Supplementary Figure 9.*** *Box plots of sensory properties, colored by sample, illustrating substantial variation among judges.*
